# Rapid Loss of CD4 T Cells by Pyroptosis During Acute SIV Infection in Rhesus Macaques

**DOI:** 10.1101/2022.05.24.493358

**Authors:** Xuan He, Malika Aid, John D. Ventura, Erica Borducchi, Michelle Lifton, Jinyan Liu, Dan H. Barouch

## Abstract

The mechanisms underlying depletion of CD4 T cells during acute HIV-1 infection are not well understood. Here we show that caspase-1-induced pyroptosis, a highly inflammatory programmed cell death pathway, is the dominant mechanism responsible for the rapid depletion of CD4 T cells in gut-associated lymphatic tissue (GALT), spleen, and lymph nodes during acute simian immunodeficiency virus (SIV) infection in rhesus macaques. Upregulation of interferon-gamma inducible factor 16 (IFI16), a host DNA sensor that triggers pyroptosis, was also observed in tissue-resident CD4 T cells and correlated with viral loads and CD4 T cell loss. In contrast, caspase-3-mediated apoptosis and viral cytotoxicity only accounted for a small fraction of CD4 T cell death. Other programmed cell death mechanisms, including mitochondria-induced caspase-independent cell death, necroptosis, and autophagy, did not significantly contribute to CD4 T cell depletion. These data support a model in which caspase-1-mediated pyroptosis is the principal mechanism that results in CD4 T cell loss in the GALT and lymphoid organs and release of proinflammatory cytokines. These findings contribute to our understanding of the pathogenesis of acute SIV infection and have important implications for the development of therapeutic strategies.

**Importance:** Different mechanisms for CD4 T cell depletion during acute HIV-1 infection have been proposed. In this study, we should in SIV infected rhesus macaques that depletion of CD4 T cells is primarily due to pyroptosis. Other pathophysiologic mechanisms may also contribute in a minor way to CD4 T cell depletion.

## Introduction

The precise mechanisms underlying CD4 T cell depletion in acute HIV-1 infection remain unknown. Previous studies have suggested that apoptosis is the main driver of CD4 T cell death during HIV-1 infection using *ex vivo* models (1, 2). However, direct infection of *ex vivo* generated susceptible cells, such as pre-stimulated CD4 T cells or immortalized T cell lines, may bias outcomes in favor of the direct viral cytopathic effects that have been known to induce apoptosis in infected cells (3–7). In contrast, a study using an *ex vivo* human lymphoid aggregate culture (HLAC) system infected with a limited number of HIV virions suggested that pyroptosis, a highly inflammatory form of programmed cell death, was the predominant mode of CD4 T cell death (8). In this study, the majority of CD4 T cell death (∼95%) occurred in uninfected bystander cells, and pyroptosis was induced via innate immune sensing of incomplete reverse transcription products accumulated during abortive HIV-1 infection. Additional evidence suggested that cell-to-cell transmission of HIV-1 was critical for activating the pyroptotic pathway, suggesting that infected CD4 T cells rather than free virions were responsible for triggering massive CD4 T cell loss (9). Autophagy, a pathway of cellular degradation independent of caspase-induced cell death, was also reported to occur within bystander CD4 T cells during *ex vivo* HIV-1 infection, contributing to CD4 T cell depletion (10).

The mechanism of systemic CD4 T cell depletion during acute HIV-1 infection *in vivo* remains to be determined. To address this, we evaluated CD4 T cell depletion in peripheral blood, spleen, pelvic lymph nodes (LN), as well as in colon and mesenteric LNs as representatives of gut-associated lymphatic tissue (GALT), which is a major site of viral replication and CD4 T cell depletion after SIV infection (4, 7). By both flow cytometric and transcriptomic analyses of CD4 T cells, we demonstrate that pyroptosis is the predominant mechanism associated with CD4 T cell depletion in peripheral blood, GALT and lymphoid organs during acute SIV infection. Such knowledge contributes to understanding SIV immunopathogenesis and may contribute to the development of novel therapies to improve immunological recovery in HIV-infected patients (11–13).

## Results

### Pathogenic SIV infection causes rapid loss of CD4 T cells in multiple tissues

To study CD4 T cell dynamics during acute and chronic SIV infection, we conducted a longitudinal study in 7 rhesus monkeys infected with SIVmac251. In peripheral blood, median peak plasma viremia occurred on day 14 following SIVmac251 infection, and high-level setpoint viremia persisted in chronic SIV infection as expected (**Figure 1A**). To evaluate cellular death pathways occurring within tissue site, we also performed serial necropsies in 19 monkeys at different times following infection, and harvested colon, mesenteric LNs, spleen and pelvic LNs on day 0 (n=3), day 3 (n=3), day 7 (n=9) and day 10 (n=4) following infection. A nested RT-PCR assay was utilized to quantitate tissue viral RNA levels as described previously (14). All tissues from day 3 following infection were negative for viral RNA (**Figure 1A**). In animals necropsied on day 7, we observed wide variations in the levels of viral RNA, about half of which showed detected levels of viral RNA. In contrast, all 4 animals necropsied on day 10 had high levels of viral RNA ranging from 1 x 10^7^ to 1 x 10^10^ RNA copies per 10^8^ cell equivalents in colon, mesenteric LNs, spleen and pelvic LNs **(Figure 1A)**. Viral DNA also showed progressive increase in tissues at necropsy from day 3 to day 10 (**Figure S1**).

**Figure 1.**
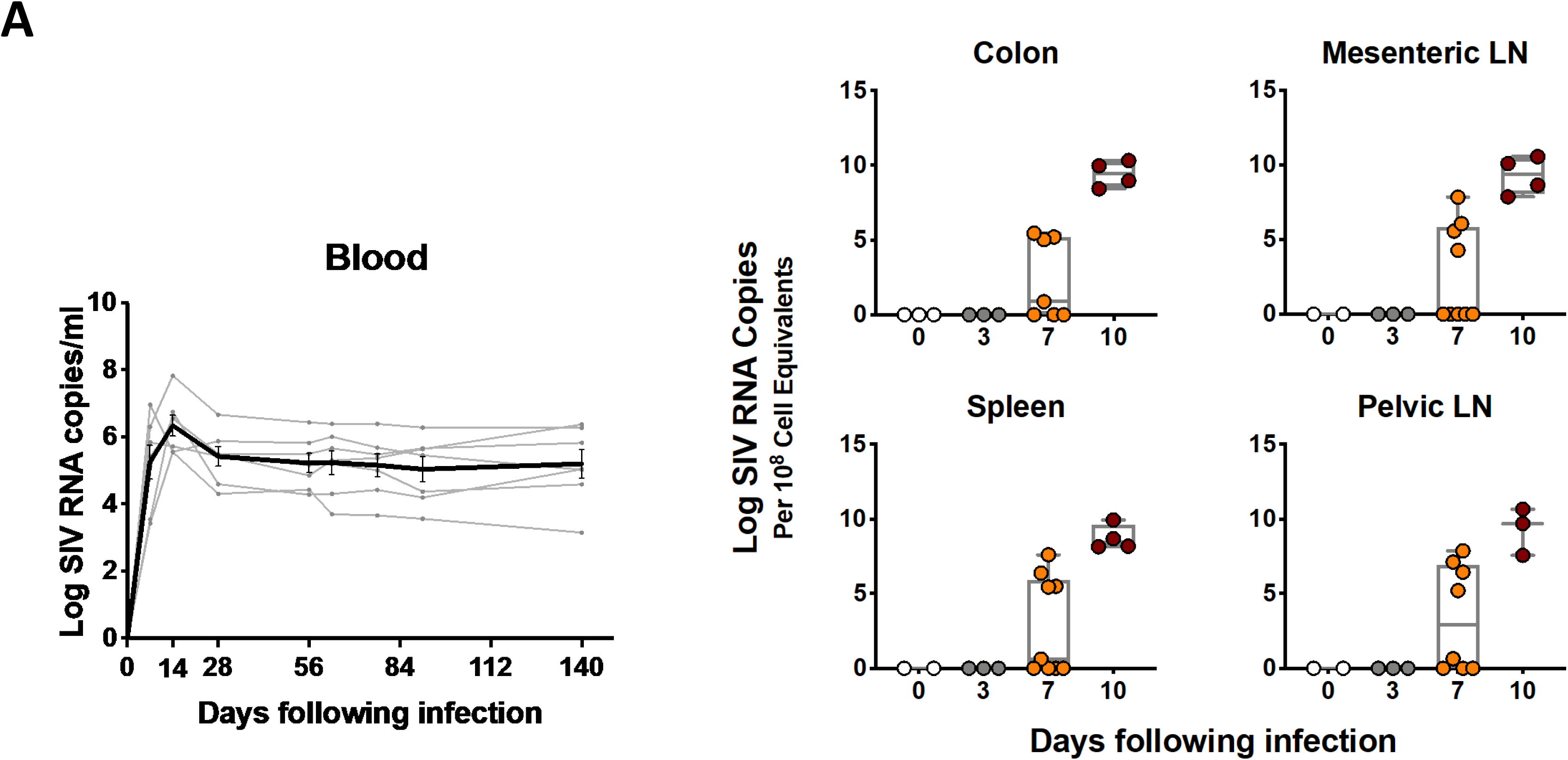
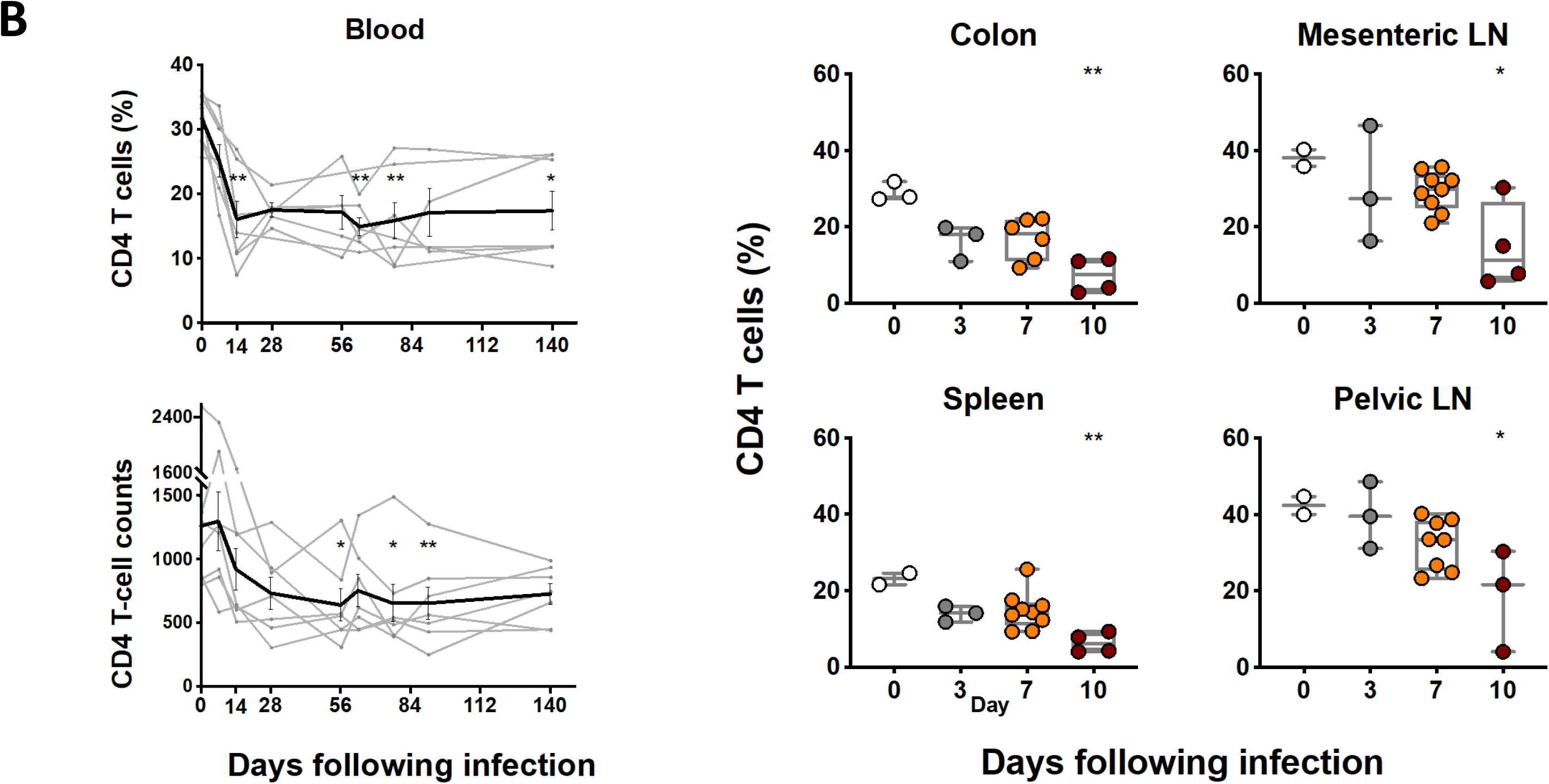
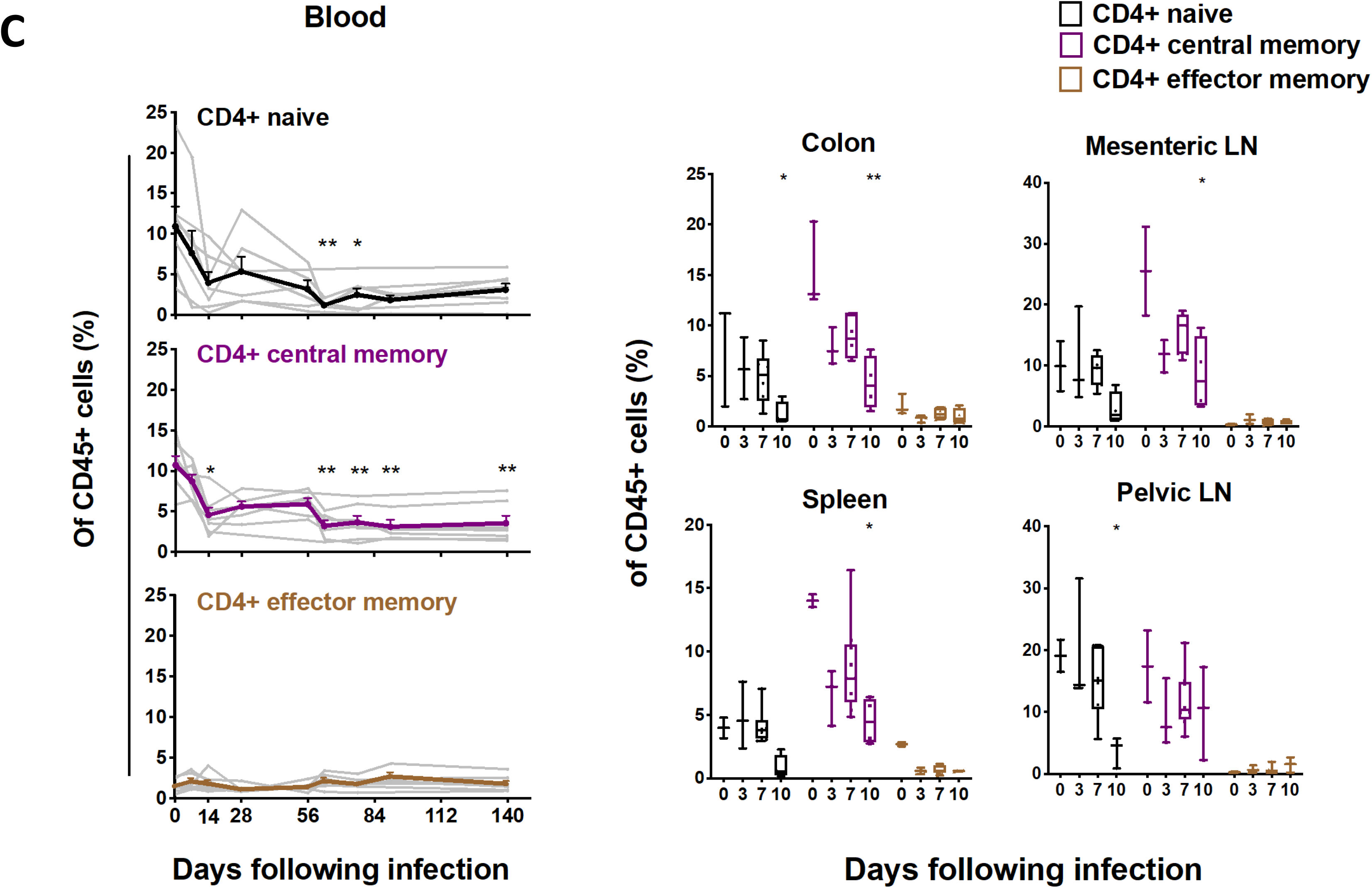
Dynamics of viral RNA and CD4 T cells in blood and tissues. (**A**) Viral RNA (log RNA copies/ml) in blood in monkeys from day 0 to 140 following inoculation with SIVmac251, and tissue viral RNA (log RNA copies/10^8^ cell equivalents) in multiple tissues at necropsies in monkeys on days 0, 3, 7 or 10 following infection with SIVmac251. (**B**) Percentage and absolute number of CD4 T cells in blood and in multiple tissues at necropsy in monkeys with SIVmac251 infection at different time points. (**C**) The dynamics of CD4 T cell naive and memory subsets in blood and different tissues during pathogenic SIVmac251 infection. Black, purple and brown correspond to naive, central memory and effector memory CD4 T cell subset, respectively. Significant *p* values are shown in figures. Experimental variables were analyzed by one-way analysis of variance (ANOVA). *\* p .05; ** p .01*.

We next evaluated the decline of CD4 T cells in blood and tissues during acute SIVmac251 infection. A significant decline of CD4 T cells was observed in peripheral blood by day 14, based on the percentage of CD4 T cells in CD45^+^ leukocytes (p=0.004) (**Figure 1B**). The low level of CD4 T cells persisted throughout chronic SIV infection. The absolute number of CD4 T cells in blood also declined progressively in acute SIV infection, with significant decrease on day 56 following SIV challenge (p=0.012) (**Figure 1B**). In tissues, the percentage of CD4 T cells in CD45^+^ leukocytes significantly dropped from a mean level of 29.0% on day 0 to 7.4% on day 10 post infection in colon (p=0.005), from 38.1% to 14.7% in mesenteric LNs (p=0.024), from 23.1% to 6.4% in spleen (p=0.008), and from 42.4% to 18.7% in pelvic LNs (p=0.034) (**Figure 1B**). This observation of dramatic loss of CD4 T cells in multiple tissues during acute SIV infection is consistent with previous studies (4, 15).

We next assessed CD4 T cell subpopulations, including naïve (CD28^+^/CD95^-^), central memory (CD28^+^/CD95^+^) and effector memory (CD28^-^/CD95^+^) cells based on CD28 and CD95 markers as described previously (15). A significant drop of central memory (CM) CD4 T cells was observed in peripheral blood by day 14 following SIV infection (p=0.02), and naïve CD4 T cells were reduced by day 63 (p=0.001) (**Figure 1C**). In tissues, the frequency of naive and CM CD4 T cells also declined significantly by day 10 following SIV infection compared to uninfected controls (For naïve CD4 in colon, p=0.048; for CM CD4 in colon, p=0.004; for CM CD4 in mLN, p=0.038; for CM CD4 in spleen, p=0.02; for naive CD4 in pLN, p=0.042) (**Figure 1C**). Additionally, we evaluated the immune status of CD4 T cells in peripheral blood and tissues. We observed an early increase of CD69^+^ CD4 T cells in colon on day 10 following challenge (p=0.04) (**Figure S2A**). In addition, we also observed a significant increase in CD4 T cell proliferation in colon (p=0.032) and in pelvic LNs (p=0.047) by days 7 and 10, respectively (**Figure S2B**), suggesting active immunological engagement by CD4 T cells during acute infection. Taken together, this longitudinal study of CD4 T cell dynamics confirmed rapid loss of CD4 T cells in both peripheral blood and tissues following pathogenic SIVmac251 infection.

### CD4 T cell depletion in GALT and secondary lymphoid tissues is strongly associated with pyroptosis

We next evaluated the potential mechanisms of CD4 T cell depletion. Activated caspase 1, a marker commonly used for pyroptosis (8), was detected longitudinally in peripheral blood and tissues. Peripheral blood CD4 T cells showed a modest increase of activated caspase 1 during acute SIV infection, which peaked on day 14 to a mean level of 5.5% following infection (p=0.03 compared to day 0) (**Figure 2A; S3A**). In contrast, there was a dramatic increase of CD4 T cell pyroptosis observed on day 10 following challenge in multiple tissues, including a mean level of 25.1% in colon (p=0.007), 10.6% in mesenteric LNs (p=0.01), 34.6% in spleen (p=0.01), and 17.6% in pelvic LNs (p=0.01) (**Figure 2B; S3B**).

**Figure 2.**
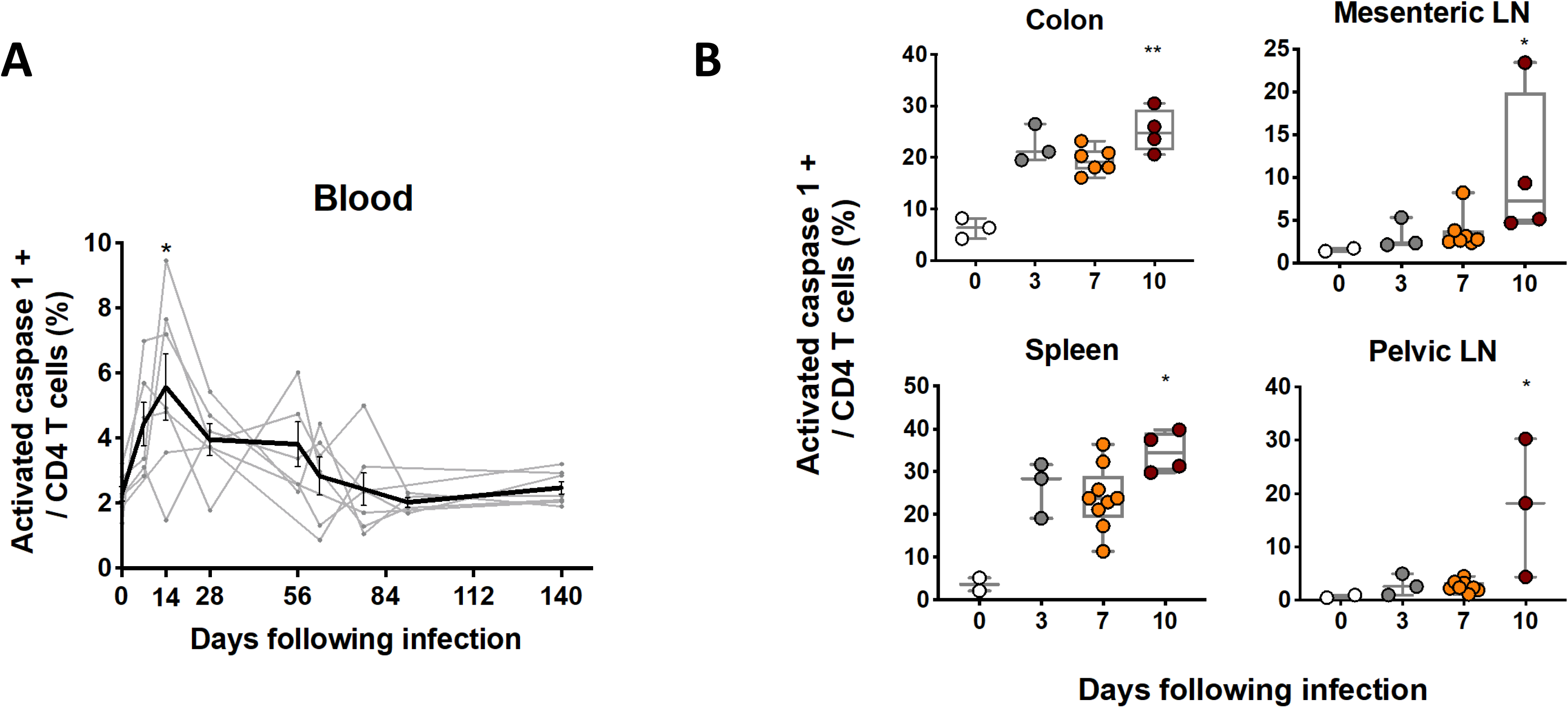
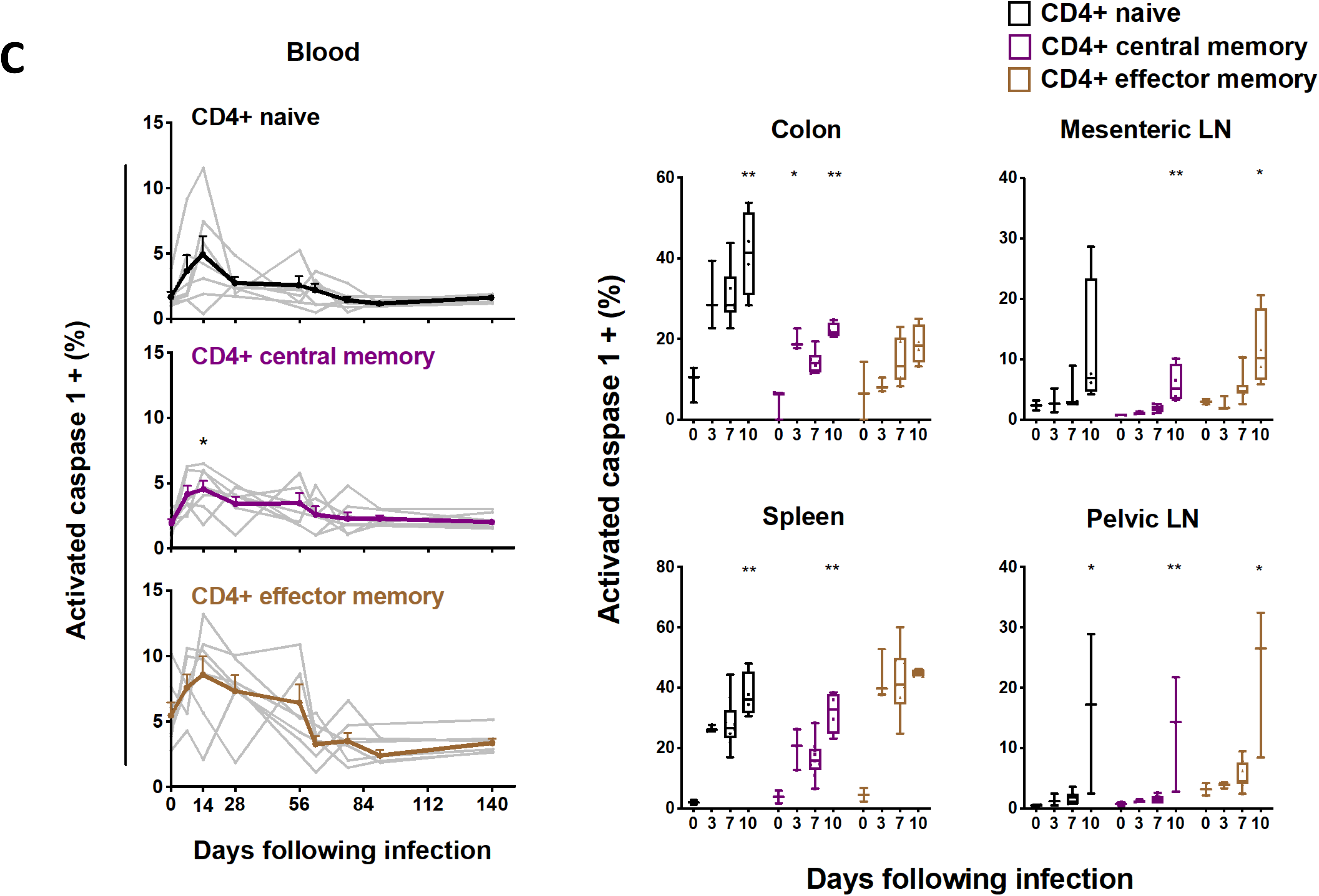
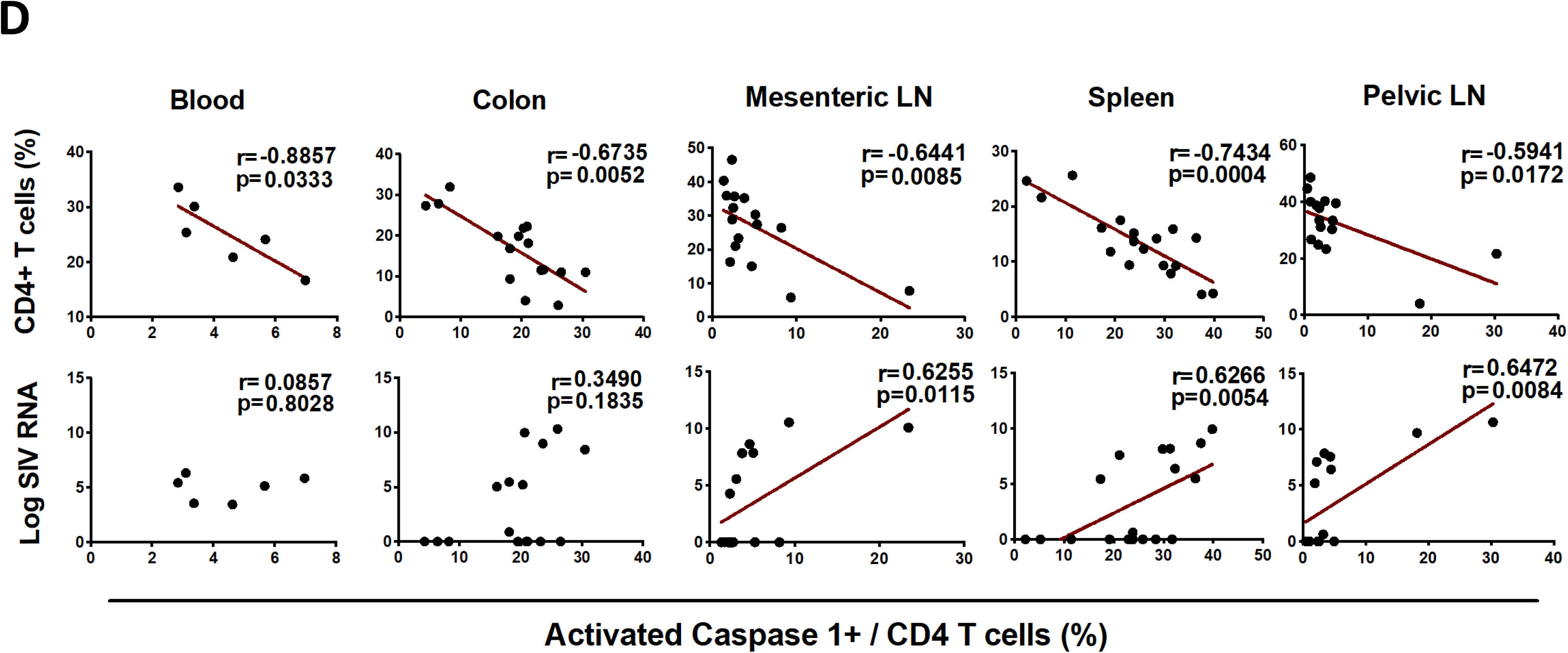
Pyroptosis of CD4 T cells in blood and tissues following SIV infection. (**A**) The line chart shows pyroptosis of CD4 T cells in blood from seven monkeys at different time points ranging from day 0 to day 140 following SIVmac251 infection. (**B**) Pyroptosis of CD4 T cells was analyzed by activated caspase 1 staining of CD4 T cells in GALT and lymphoid tissues at necropsy during pathogenic SIVmac251 infection. (**C**) Pyroptosis of naive and memory CD4 T cell subsets in blood and different tissues at necropsy during pathogenic SIVmac251 infection. Black, purple and brown correspond to naive, central memory and effector memory CD4 T cell subset, respectively. (**D**) The correlation between the expression of activated caspase 1 in CD4 T cells and the percentage of CD4 T cells or the magnitude of viral RNA in blood on day7 and in multiple tissues from all the monkeys at necropsy. Solid lines indicate the correlation with significant *p* values. Significant *p* values are shown in figures. Experimental variables were analyzed by one-way analysis of variance (ANOVA). *\* p .05; ** p .01.* For correlation analysis, spearman rank order correlation coefficients r and corresponding p values are indicated.

We also assessed pyroptosis of CD4 T cells by evaluating the magnitude of activated caspase 1 in CD4 naive and memory subsets in blood and tissues. Generally, lower expression of caspase 1 was found in CD4 naive and memory cells in blood compared to tissues (**Figure 2C**). Increased pyroptotic activity was observed in CD4 naive subset in most tissues as early as day 10 (For colon, p=0.009; for spleen, p=0.008; for pLN, p=0.013). CD4 CM cells showed increased pyroptosis in blood, GALT and lymphoid tissues on day10 following infection (For colon, p=0.003; for mLN, p=0.004; for spleen, p=0.005; for pLN, p=0.008). Although we did not detect an obvious decline in CD4 effector memory (EM) cells, which are a small subpopulation of CD4 in blood and tissues, an increase in pyroptotic activity was observed in EM CD4 cells, reaching significance in mLN (p=0.045) and pLN (p=0.03) during early SIV infection **(Figure 2C)**.

We next assessed whether CD4 T cell pyroptosis was associated with CD4 T cell depletion and viral RNA in blood and tissues during SIV infection. CD4 T cell pyroptosis showed a strong negative correlation with the percentage of CD4 T cells from peripheral blood in rhesus monkeys on day 7 following infection (r=-0.88, p=0.03) (**Figure 2D**). Similarly, negative correlations were observed between pyroptosis and CD4 T cell level in all tissues evaluated, including colon (r=-0.67, p=0.005), mesenteric LNs (r=-0.64, p=0.008), spleen (r=-0.74, p=0.0004) and pelvic LNs (r=-0.59, p=0.01). CD4 T cell pyroptosis was also positively correlated with viral RNA in spleen (r= 0.62, p=0.005), mesenteric LNs (r=0.62, p=0.01) and pelvic LNs (r=0.64, p=0.008) (**Figure 2D**). Additionally, the level of naive CD4 T cells showed significantly negative correlations with its pyroptotic activity in colon (r=-0.66, p=0.004) and pelvic LNs (r=-0.56, p=0.02), and percentage of CM CD4 T cells was also highly associated with its pyroptosis in colon (r=-0.76, p=0.0006), mesenteric LNs (r=-0.59, p=0.01) and spleen (r=-0.76, p=0.0002) (**Figure S3C**). Taken together, these data suggest that pyroptosis was highly associated with reduced CD4 T cells, including naïve and central memory CD4 T cells, and was correlated with levels of viral RNA in lymphoid organs.

### SIV-mediated apoptosis was only minimally detected in CD4 T cells during acute SIV infection

Multiple reports have suggested increased apoptosis of CD4 T cells during SIV infection (4, 16, 17). To evaluate the contribution of apoptotic activity to CD4 T cell death, we investigated the kinetics of CD4 T cell apoptosis via flow-cytometric detection of intracellular activated caspase 3, a well-characterized effector marker of apoptosis (8, 16). There was a detectable but marginal increase of CD4 T cells expressing caspase 3 on day 14 compared to day 0 in peripheral blood during early SIV infection (p=0.008) (**Figure 3A; S4A**). In tissues, modest increases of caspase 3 expression in CD4 T cells were also observed by day 10 following SIVmac251 infection in colon (p=0.04) and spleen (p=0.03) with less than 4% tissue-resident CD4 T cells undergoing apoptosis (**Figure 3B; S4B**). No obvious change of CD4 T cell apoptosis was observed in CD4 T cell naive or memory subsets during early SIV infection (**Figure S4C**). CD4 T cell apoptosis was negatively correlated with percentage of CD4 T cells from colon (r=-0.60, p=0.01) and mesenteric LNs (r=-0.63, p=0.004) (**Figure 3C**). Notably, in contrast with caspase-1-mediated pyroptosis, caspase-3-mediated apoptosis was positively correlated with viral RNA in colon (r=0.68, p=0.004). Caspase-3-mediated apoptosis accounted for <4% of CD4 T cell death in GALT and lymphoid organs.

**Figure 3.**
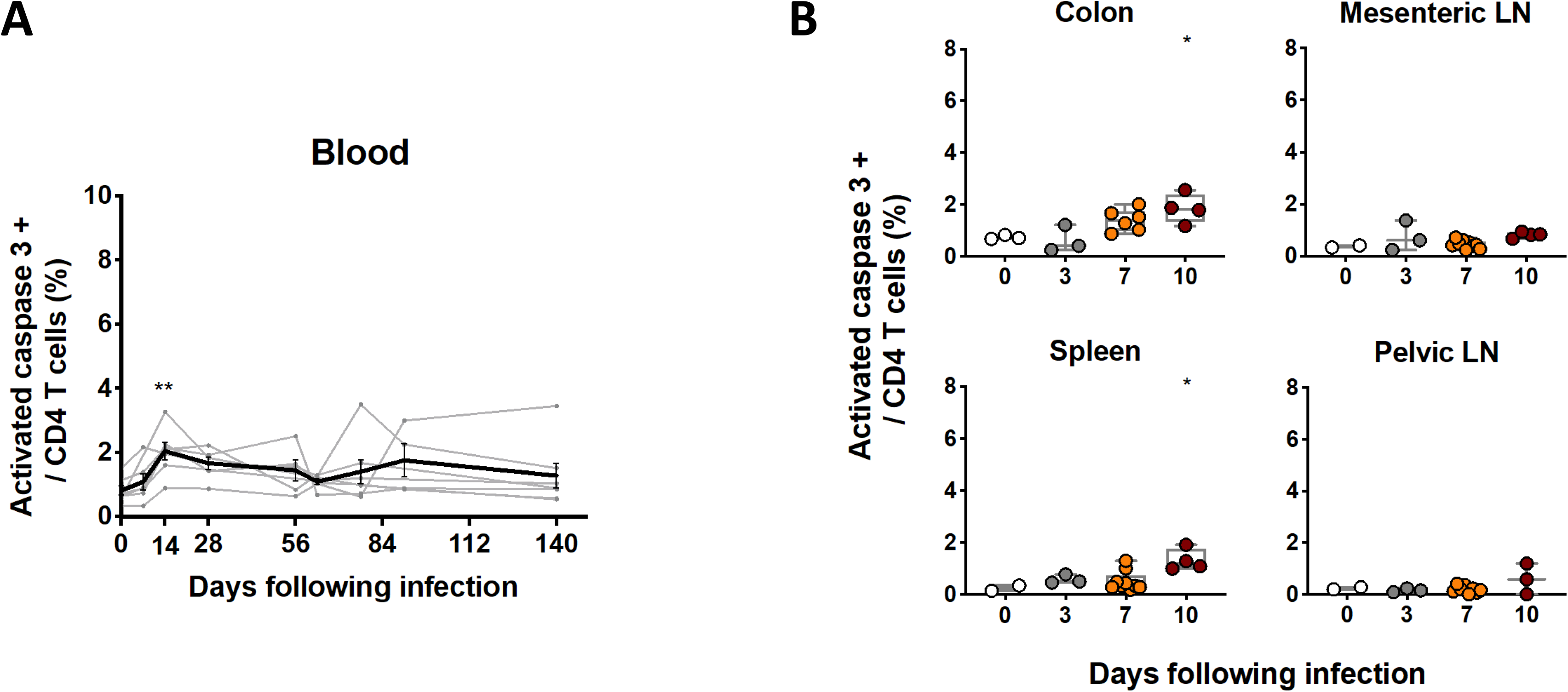
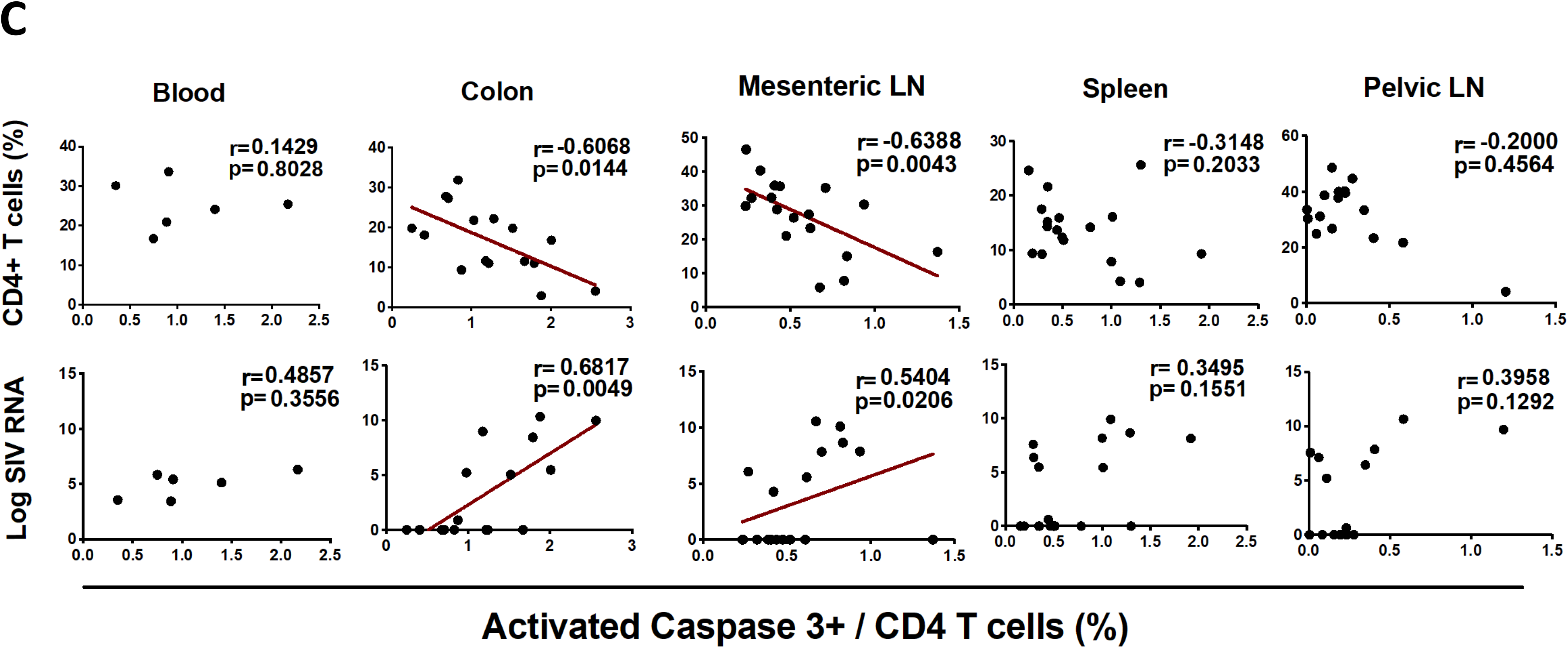
Apoptosis of CD4 T cells in blood and tissues following SIV infection. (**A**) The line chart shows apoptosis of CD4 T cells in blood from seven monkeys at different time points ranging from day 0 to day 140 following SIVmac251 infection. (**B**) Apoptosis of CD4 T cells was analyzed by activated caspase 3 staining of CD4 T cells in multiple tissues at necropsy during pathogenic SIVmac251 infection. (**C**) The correlation between the expression of activated caspase 3 in CD4 T cells and the percentage of CD4 T cells or the magnitude of viral RNA in blood on day 7 and in multiple tissues from all the monkeys at necropsy. Solid lines indicate the correlation with significant *p* values. Significant *p* values are shown in figures. Experimental variables were analyzed by one-way analysis of variance (ANOVA). *\* p .05; ** p .01.* For correlation analysis, spearman rank order correlation coefficients r and corresponding p values are indicated.

### Other programmed cell death mechanisms contribute minimally to CD4 T cell depletion

We next performed a longitudinal transcriptomic analysis of sorted CD4 T cells in mesenteric and pelvic lymph nodes between day 0 and day 10 via bulk RNA-seq. We observed a significant increase of the number of reads mapped to the SIV genome on day 10 in total RNA extracted from CD4 T cells in LNs (p<0.05) (**Figure S5**), indicating the presence of productive SIV within CD4 T cells following infection. The number of reads mapped to SIV were also associated with viral loads (r=0.66, p=0.00005) and CD4 T cells (r=-0.58, p=0.00056) as expected (**Figure S5**). To evaluate the direct impact of SIVmac251 infection on CD4 T cell death, we quantified the percentage of CD4 T cells containing SIV transcripts. A flow cytometry-based assay capable of measuring SIV transcription in CD4 T cells undergoing infection was utilized. During early SIV infection in monkeys, less than 0.5% of CD4 T cells in peripheral blood exhibited productive SIV infection (**Figure 4A)**. Higher frequencies of SIV+ CD4 T cells were observed in lymphoid tissues on day 10 following SIV infection, but still <5% of CD4 T cells **(Figure 4A)**. As expected, SIV-infected CD4 T cells from blood or tissues resided in CD95^+^ memory subsets with high levels of CCR5 expression **(Figure S6)**. Considering that more than 95% of CD4 T cells were negative for SIV-associated transcripts and yet we detected on average 10∼35% of the entire CD4 T cell population expressing activated caspase 1 in tissues, it suggested that the vast majority of caspase-1-expressing pyroptotic cells during acute SIV infection were non-productively infected bystander cells.

**Figure 4.**
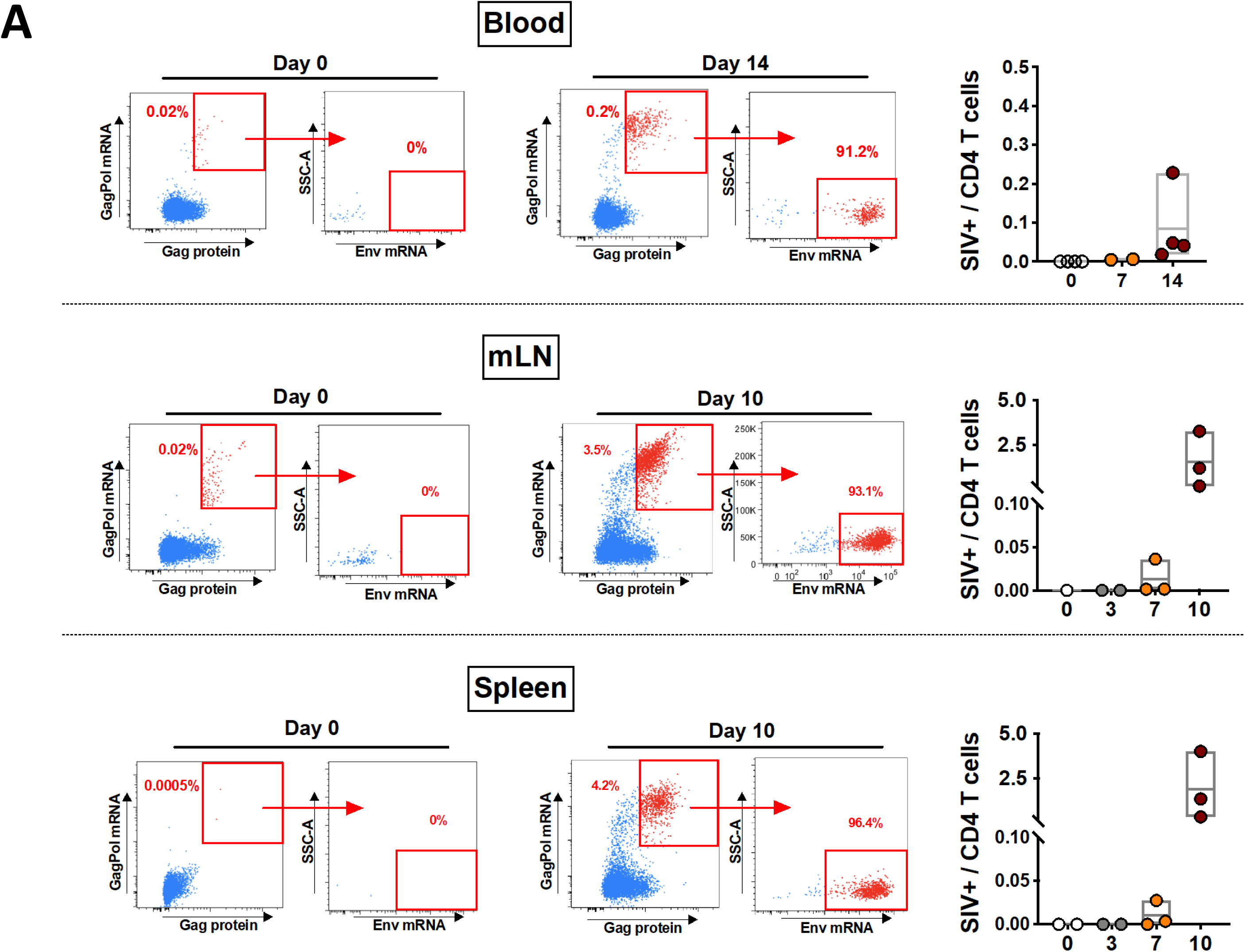
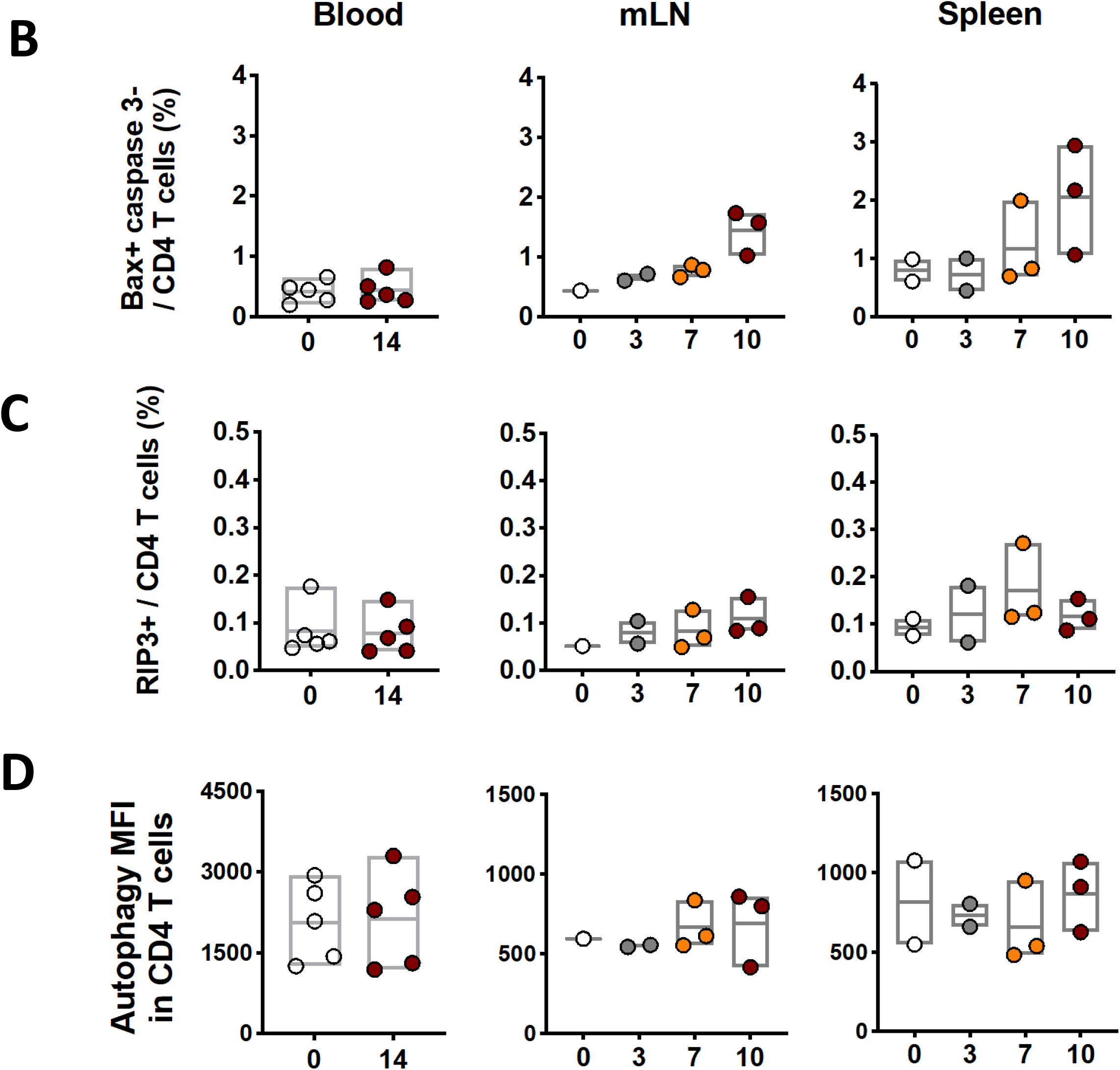
Quantification of SIV-infected CD4 T cells with translation-competent virus, and mitochondria-induced caspase-independent cell death, necroptosis or autophagy of CD4 T cells in blood and tissues. (**A**) Representative flow cytometric plots showing CD4 T cells with or without translation-competent SIV in blood on day 0 or day 14 following infection, as well as in mesenteric LNs and spleen on day 0 or day 10 following infection. CD4 T cells with productive SIV infection were quantified in blood on day 0 and 14, and in mLN and spleen on day 0, 3, 7, or 10 during SIVmac251 infection. (**B**) CD4 T cells undergoing mitochondria-induced caspase-independent cell death were analyzed in blood and tissues during SIVmac251 infection. (**C**) CD4 T cells undergoing necroptosis were analyzed in blood and tissues during SIVmac251 infection. (**D**) CD4 T cells undergoing autophagy were analyzed in blood and tissues during SIVmac251 infection.

Caspase-independent cell death (CICD) is an apoptotic pathway that does not include executioner caspase activation (e.g. caspase 1, 3, and 11) (18–20). HIV-1 Vpr has been reported to kill cells by this pathway, in which mitochondrial membrane permeabilization was involved as a crucial step (21). *BCL2L14* and *BAK1*, which both belong to the *Bcl-2* gene family and are capable of regulating apoptotic pathway via mitochondria (22), were upregulated in CD4 T cells from mLNs (*BCL2L14*, p=0.01; *BAK1*, p=0.009) and pLNs (*BCL2L14*, p=0.02; *BAK1*, p=0.04) on day 10 following SIV infection by transcriptomic profiling. These data suggested activation of mitochondria in LN-resident CD4 T cells during early SIV infection. In addition, based on flow cytometric and transcriptomic analyses, we did not observe the downstream activation of executioner caspases, which suggested the possibility of mitochondria-induced CICD in CD4 T cells that may be contributing to CD4 T cell depletion during early SIV infection. We therefore checked the proportion of CD4 T cells with CICD-associated gene expression in blood or tissues from SIV-infected monkeys. Bax protein, which is essential in triggering permeabilization of mitochondria (23), was introduced in the flow cytometric analyses. As expected, CD4 T cells undergoing apoptosis expressed both Bax and caspase 3 (**Figure S7A**). Of note, we observed the CD4 T cell populations with Bax expression but negative for caspase 3 which were qualified for carrying out mitochondria-induced CICD. However, no significant changes regarding the percentage of Bax^+^ caspase 3^-^ CD4 T cells were observed in blood, spleen and mLN during the course of early infection (**Figure 4B**). In order to account for additional cell death possibilities, we assessed the role of necroptosis and autophagy during early SIV infection. Receptor interacting protein 3 (RIP3), a marker for necroptosis (24), was assessed in CD4 T cells from blood and lymphoid tissues. Less than 0.5% of CD4 T cells expressing RIP3 proteins were observed in blood or tissues, and RIP3^+^ CD4 T cells stayed at low level during early SIV infection (**Figure 4C; S7B**). Although HIV Env-mediated autophagy was found in CD4 T cells in HIV infection when apoptosis was blocked (10), we did not observe any significant changes regarding autophagy of CD4 T cell population during early SIV infection (**Figure 4D; S7C**).

### Transcriptional changes in CD4 T cells during early SIV highlight potent pyroptotic activity of CD4 T cells

To further elucidate the mechanistic basis of CD4 T cell depletion following early SIV infection, we evaluated transcriptional profiling of CD4 T cells from mesenteric and pelvic lymph nodes following SIV infection by bulk RNA-seq. Approximately 1000 genes were captured and showed significant changes in CD4 T cells from mLNs and pLNs on day 10 following infection compared to uninfected controls, of which 493 genes showed significant differential expression in both mLNs and pLNs (**Figure 5A**). Gene set enrichment analysis (GSEA) revealed significant enhancement of interferon (mLNs, NES=3.13, FDR q value< 10^-6^; pLNs, NES=3.21, FDR q value <10^-6^) and inflammatory response pathways (mLNs, NES=2.63, FDR q value < 10^-6^; pLNs, NES=2.72, FDR q value <10^-6^) in CD4 T cells on day 10 following infection compared to uninfected controls (**Figure 5B**). Transcriptomic analyses showed further evidence of SIV replication in CD4 T cells, as shown by upregulation of HIV infection pathways (mLNs, NES=1.38, FDR q value=0.02; pLNs, NES=1.55, FDR q value=0.02) (**Figure 5B**).

**Figure 5.**
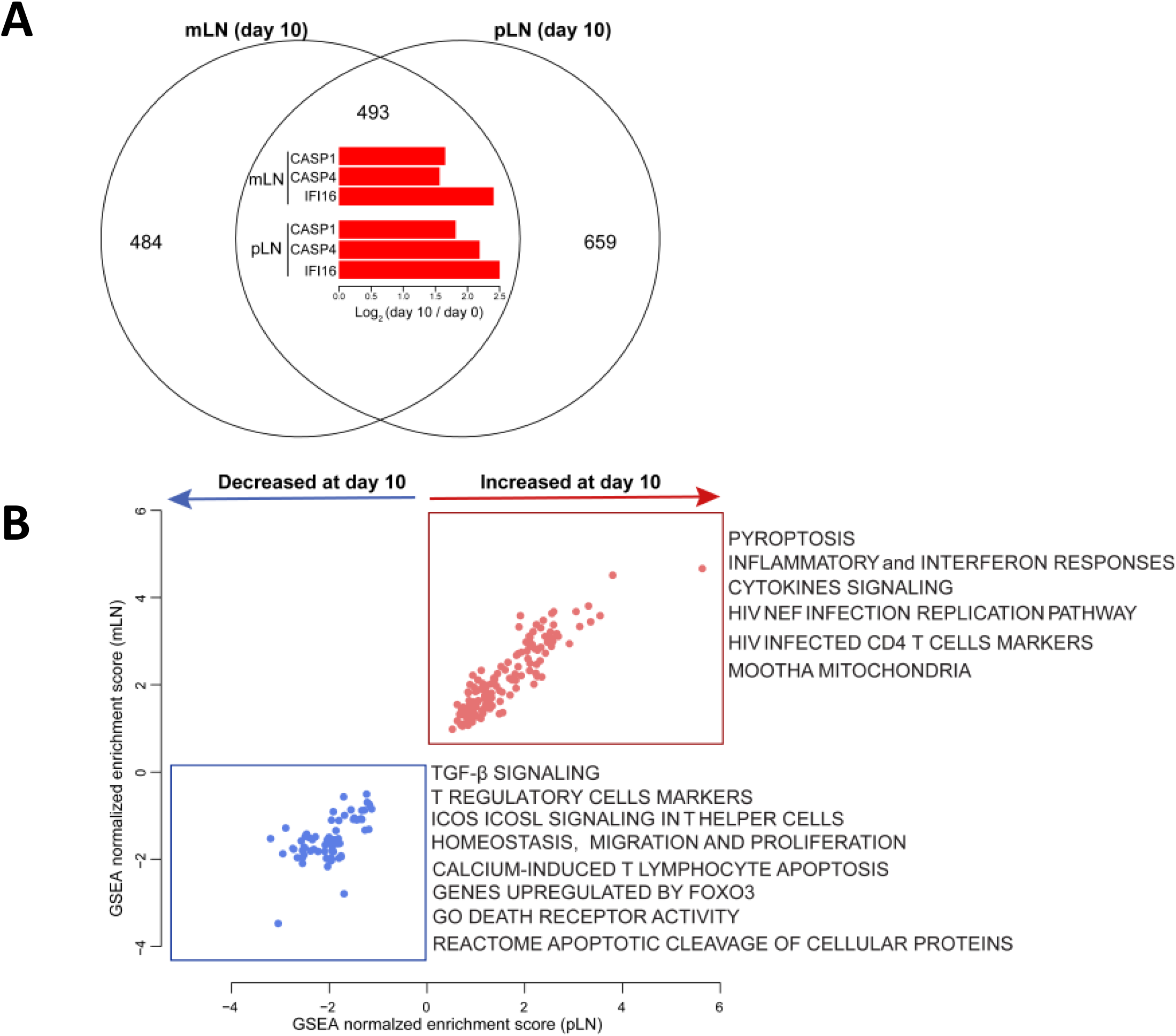
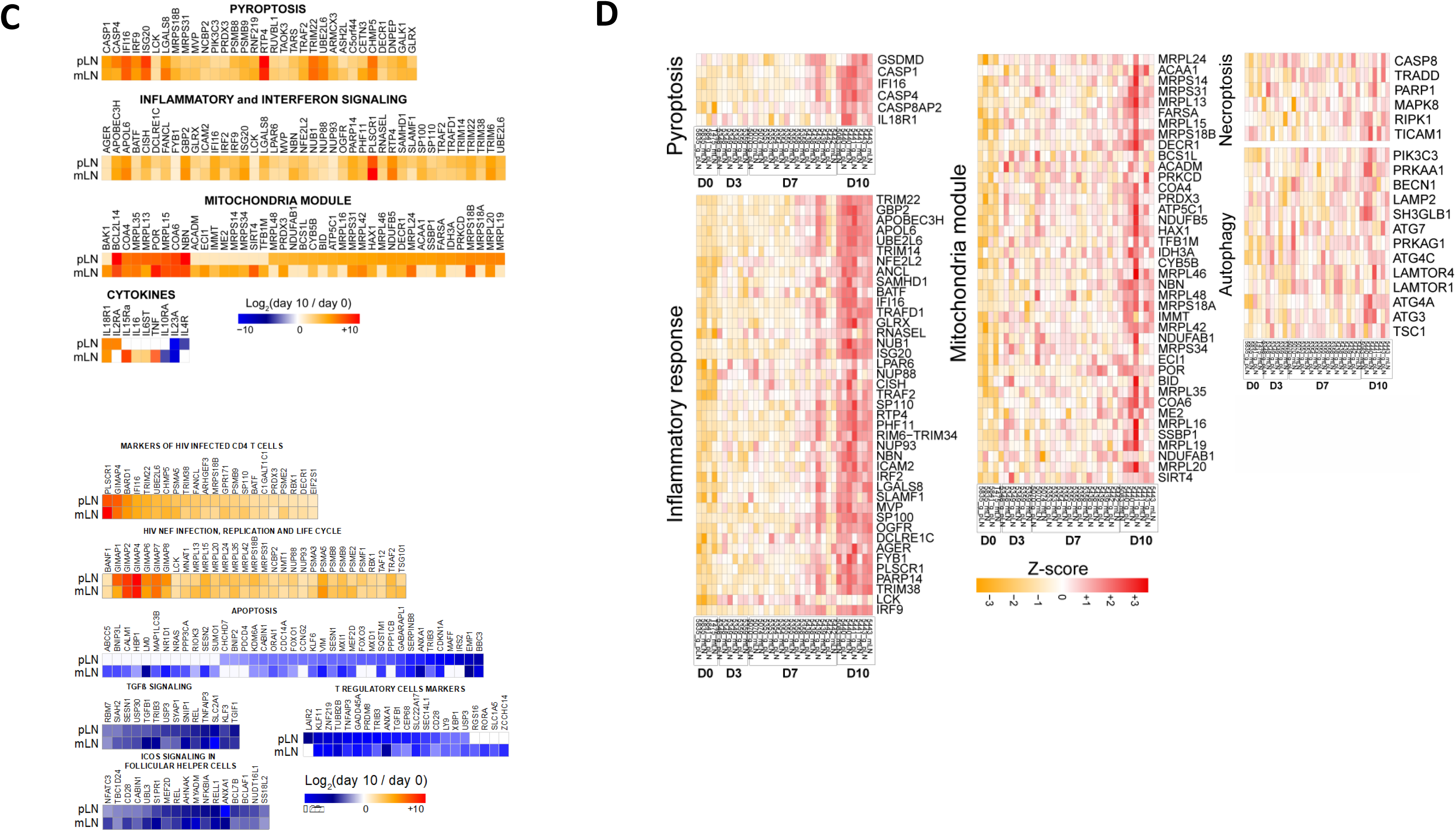
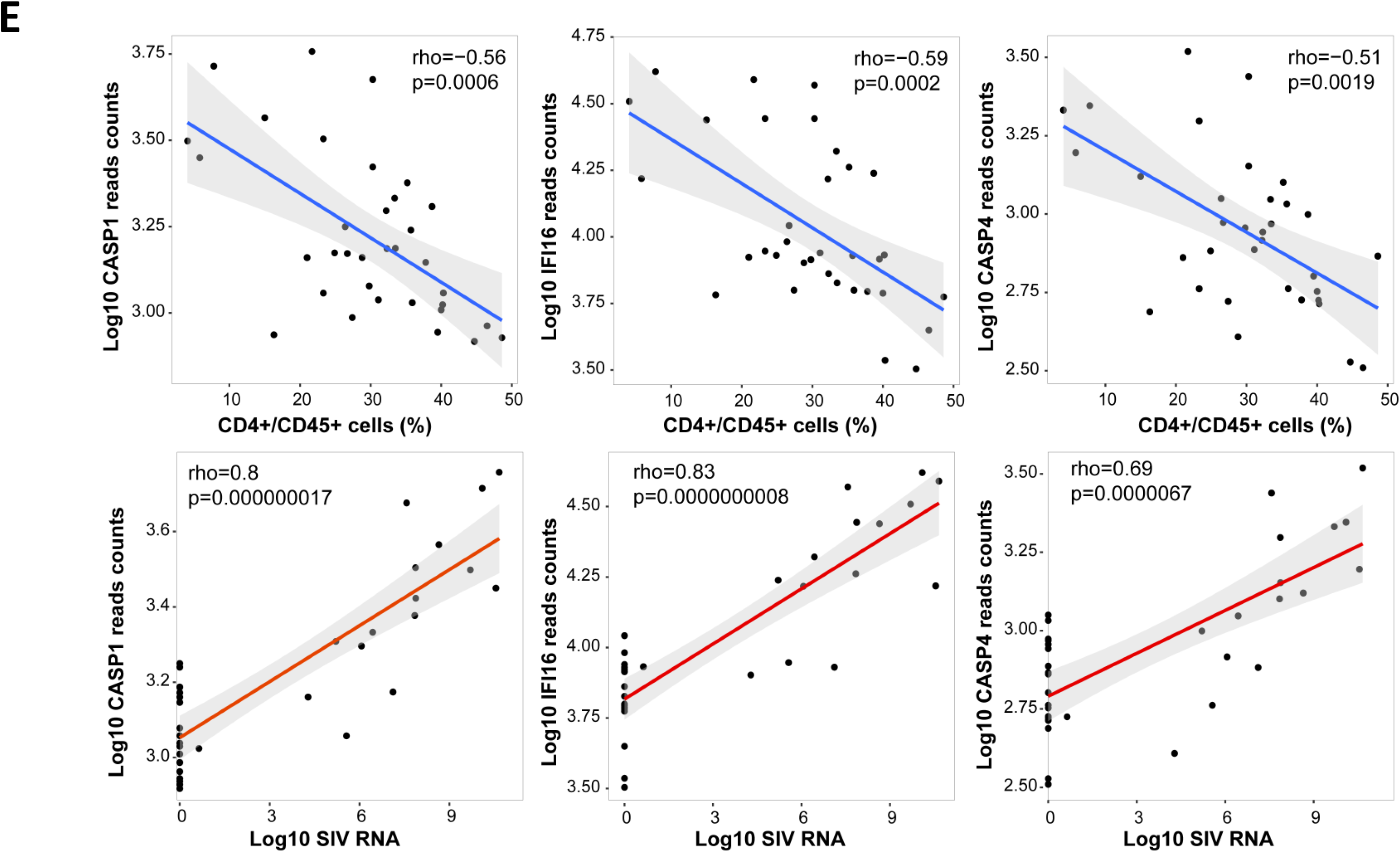
Pyroptosis and inflammatory signaling in sorted CD4 T cells from mLN and pLN from SIV-infected monkeys. (**A**) Venn diagram (top) of genes significantly increased or decreased at day 10 compared to day 0 in sorted CD4 T cells from mLN and pLN in SIV infected monkeys. All genes with a BH-corrected p of 0.05 were selected (493 common genes, 484 genes unique to mLN and 659 unique to pLN). The common plot area exhibited barplots of the log2 fold change expression (day10/day 0) of pyroptotic markers *CASP1*, *CAPS4* and I*FI16* in mLN and pLN. (**B**) The scatter plots represent the normalized enrichment score (NES) of the top enriched pathways at day 10 compared to day 0 in mLN and pLN. Pathways increased at day 10 were represented in red and pathways decreased at day 10 were represented in blue dots, where each dot corresponds to an individual pathway. X-axis represents the pathways NES in pLN and y-axis the pathways NES in mLN. Each dot on the plot represents a pathway, where increased pathways are represented in red dots and decreased pathways are represented in blue dots. (**C**) Heatmaps showing the log_2_ fold change (day 10 / day 0) of the significant genes in pathways shown in panel B. Color gradient represent the log2 fold change ranging from blue (decreased) to white (not significant) to red (increased). All genes were selected using a BH-corrected p of 0.05. (**D**) Heatmaps of the row scaled normalized-expression of markers of pyroptosis, inflammatory response, mitochondria module, necroptosis, autophagy. Each column represents an individual animal separated by time points. Each row represents an individual gene. Heatmap color gradient represent the Z-score scaled expression of genes at days 0, 3, 7 and 10, ranging from yellow (decreased) to white (not significant) to red (increased). Animals are grouped by time points as follow: baseline (day 0) and following infection (days 3, 7, 10). (**E**) Scatter plots of the log transformed expression of pyroptotic markers *CASP1*, *CASP4*, *IFI16*, correlated with the frequency of CD4 T cells (top) and viral RNA (bottom) at days 0, 3, 7 and 10 following SIV infection. Spearman correlation coefficient (ρ) and p value are shown for each gene. Gray area represents the confidence interval (IC) of 95%.

Of importance, GSEA reported an induction of pyroptotic signaling in CD4 T cells from both mLNs and pLNs on day 10 following SIV infection (**Figure 5B-D**). Consistent with the flow cytometric analysis, caspase 1 gene *CASP1* was significantly upregulated in CD4 T cells from mLNs (p=0.00002) and pLNs (p=0.0006) on day 10 compared to day 0 based on gene expression analysis. We also observed upregulation of *CASP4* in mLNs (p=0.0001) and pLNs (p=0.0004). It has been reported that caspase 4 may trigger activation of caspase 1 in macrophages during dengue infection (25). Previous reports have also identified interferon-gamma–inducible protein 16 (IFI16) as a key inflammasome DNA sensor, thus making IFI16 an important component in the canonical pyroptotic pathway (26). *IFI16* was strongly upregulated in CD4 T cells from mLNs (p=0.000009) and pLNs (p=0.001) on day 10 following SIV infection **(Figure 5C, D**). Gasdermin D (*GSDMD*), a marker of pyroptosis, was observed with trend of upregulation on day 10 (p=0,08), and was highly associated with viral loads (r=0.64, p=0.000054) in LNs. The proinflammatory cytokines *IL16* and *TNF* as well as the receptor genes for inflammatory cytokine receptors including *IL6ST*, *IL15RA*, *IL18R1* were upregulated in CD4 T cells from lymph nodes on day 10 following infection compared to uninfected controls (*IL16* in mLN, p=0.02; *TNF* in mLNs, p=0.04; *IL6ST* in mLNs, p=0.007; *IL15RA* in mLNs, p=0.01; *IL-18R1* in pLNs, p=0.03) (**Figure 5C**). We also checked genes that are relevant to other cell death pathways including apoptosis, autophagy and necroptosis in CD4 T cells from lymph nodes, and transcriptomic profiling revealed no significant upregulation of these pathways (**Figure 5C, D**).

Expression of these genes associated with pyroptosis, including *CASP1*, *CASP4 and IFI16*, were inversely correlated with CD4 T cell level (*CASP1*, r=-0.56, p=0.0006; *CASP4*, r=-0.51, p=0.019; *IFI16*, r=-0.59, p=0.0002) and were positively correlated with viral loads in these animals (*CASP1*, r=0.80, p<0.0001; *CASP4*, r=0.69, p<0.0001; *IFI16*, r= 0.83, p<0.0001) (**Figure 5E**). Genes associated with inflammatory pathways were also highly correlated with CD4 T cell frequency and viral loads in the macaques (**Figure S8**). Taken together, these data demonstrate upregulation of genes involved in the pyroptotic pathway in SIV-infected rhesus macaques, consistent with the flow cytometric studies, suggesting that pyroptosis plays a key role in CD4 T cell depletion during acute SIV infection.

## Discussion

This study extends prior *ex vivo* studies (8, 9) and provides comprehensive *in vivo* evidence that pyroptosis is the predominant mechanism of systemic CD4 T cell depletion in SIV-infected rhesus macaques. Pyroptosis of CD4 T cells was observed by both flow cytometry and transcriptomics, and was particularly evident in GALT and lymphoid tissues (4, 28).

Our results confirm and extend a previous study showing upregulation of genes involved in pyroptosis pathway in draining LNs during early SIV infection by bulk RNA-seq (27). We investigated immunologic and transcriptomic profiles of systemic CD4 T cells from GALT and lymphoid tissues during acute SIV infection and assessed the contributions from all potential programmed cell death mechanisms to CD4 T cell death. Our data suggest that the pyroptosis of CD4 T cells was the dominant mode responsible for CD4 T cell loss either in GALT or lymphoid tissues. In addition, we observed lower pyroptosis of CD4 T cells in peripheral blood compared with lymphoid tissues, consistent with a previous report showing that blood CD4 T cells are partially resistant to pyroptosis due to deeper resting state with less DNA sensor IFI16 (29). Moreover, CD4 T cells in lymphoid tissues with higher proportions of SIV infection may facilitate pyroptosis of surrounding bystander CD4 T cells by cell-to-cell interactions (9). Our transcriptomic data also highlight the potential role of IFI16, a HIV-1 DNA sensor which leads to caspase 1 activation. Our *in vivo* data suggest a critical role for IFI16, which detects viral DNA transcripts and triggers inflammasome assembly, which then leads to activation of caspase 1 and pyroptosis in CD4 T cells.

The study of HIV-1 infection using HLAC system highlighted the massive death caused by pyroptosis occurring in bystander CD4 T cells (8). During SIV infection, we observed profound depletion of naive CD4 T cells derived from tissues and associated with pyroptosis. The depletion of naive CD4 T cell populations, provides indirect evidence for a strong bystander effect, especially within the gastrointestinal tract and other tissue compartments. Moreover, the drastic loss of naive or central memory CD4 T cells causes the insufficient generation and delivery of CD4 T cell effectors fighting against SIV infection and leads to onset of progressive SIV pathogenesis.

In contrast with pyroptosis, we observed only a small proportion of CD4 T cells expressing caspase 3, indicative of minimal apoptosis of CD4 T cells in peripheral blood and tissues during early SIV infection. Our data are consistent with the *ex vivo* study using the HLAC system [8], suggesting that apoptosis may be a minor component of CD4 T cell depletion *in vivo*. Moreover, the low frequency of CD4 T cells undergoing productive SIV infection in blood and tissues indicate that direct virus cytotoxicity provides only minimal contribution to CD4 T cell loss. Another *ex vivo* study has observed that the activation of DNA-dependent protein kinase (DNA-PK) contributes to CD4 T cell death during the process of viral integration (3). In our study, no obvious change regarding gene expression of DNA-PK was observed in CD4 T cells. Moreover, other programmed cell death pathways including autophagy, necroptosis and mitochondria-induced CICD appeared to barely play roles in CD4 T cell depletion during acute SIV infection.

Taken together, our observations show that CD4 T cell depletion in acute SIV infection in rhesus macaques is primarily the result of caspase-1-mediated pyroptosis. These findings have important implications for our understanding of HIV-1 pathogenesis and may provide new therapeutic targets for immunological recovery.

## Acknowledgements

We thank Joern Schmitz, Lawrence Tartaglia, Daniel Ram, Venous Hamza, Abishek Chandrashekar, Jingyou Yu for advice, assistance and reagents. We acknowledge support from the NIH (OD024917, AI149670, AI128751, AI129797, AI126603, AI124377, AI164556) and the Ragon Institute of MGH, MIT, and Harvard.

## Authorship contributions

D.H.B., X.H. designed the study and reviewed all data. X.H. performed the immunological assays and analyzed data. M.A. performed the transcriptomic analysis. D.H.B, J.D.V. edited the manuscript. E.B., M.L., J.L. contributed to materials. X.H. and D.H.B. wrote the paper with all co-authors.

## Declaration of Interests

No competing financial interests.

## Supplemental Figure legends

**Figure S1.**
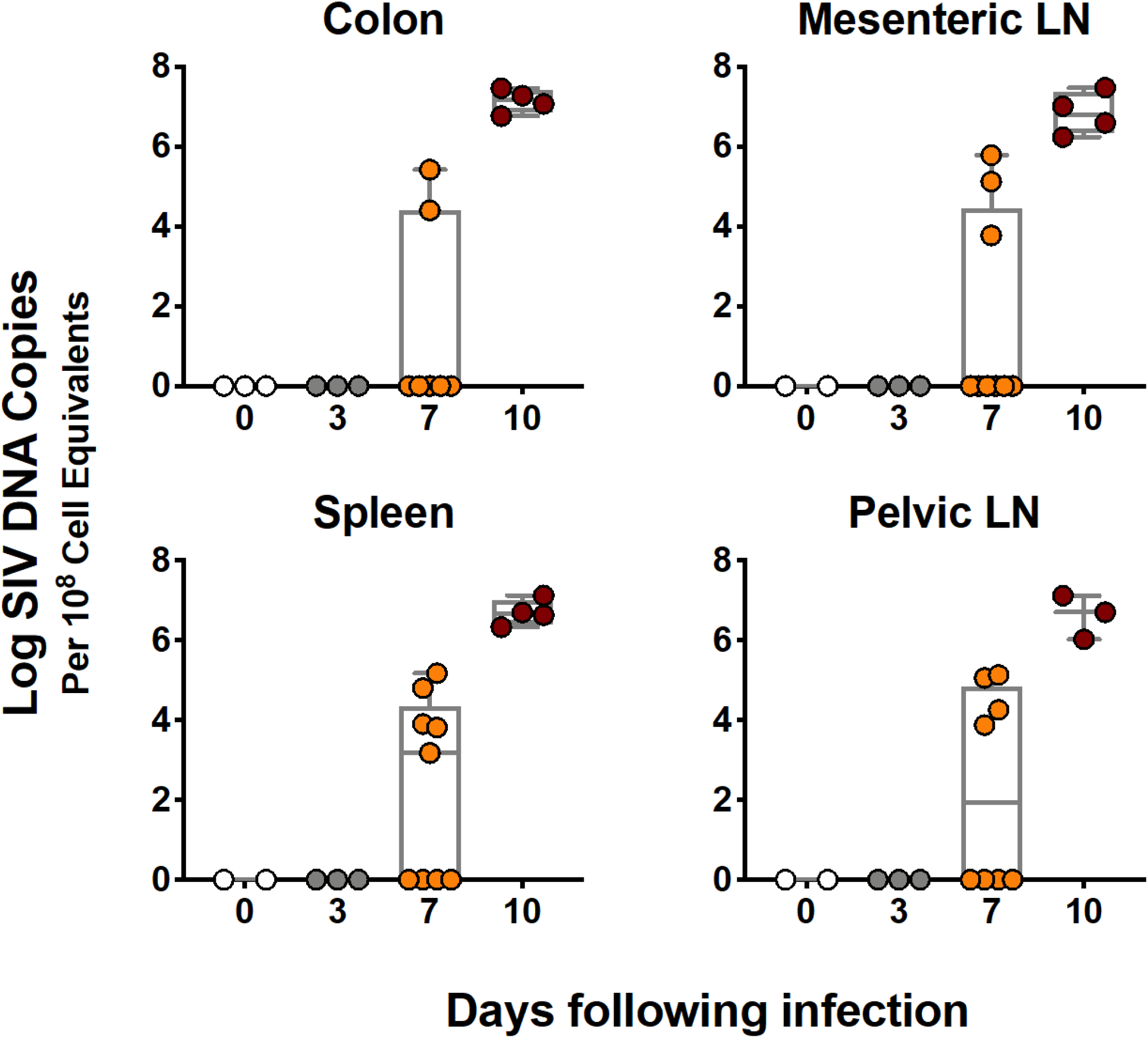
Dynamics of viral DNA in tissues. Viral DNA (log RNA copies/10^8^ cell equivalents) in multiple tissues at necropsies in monkeys on days 0, 3, 7 and 10 following infection with SIVmac251.

**Figure S2.**
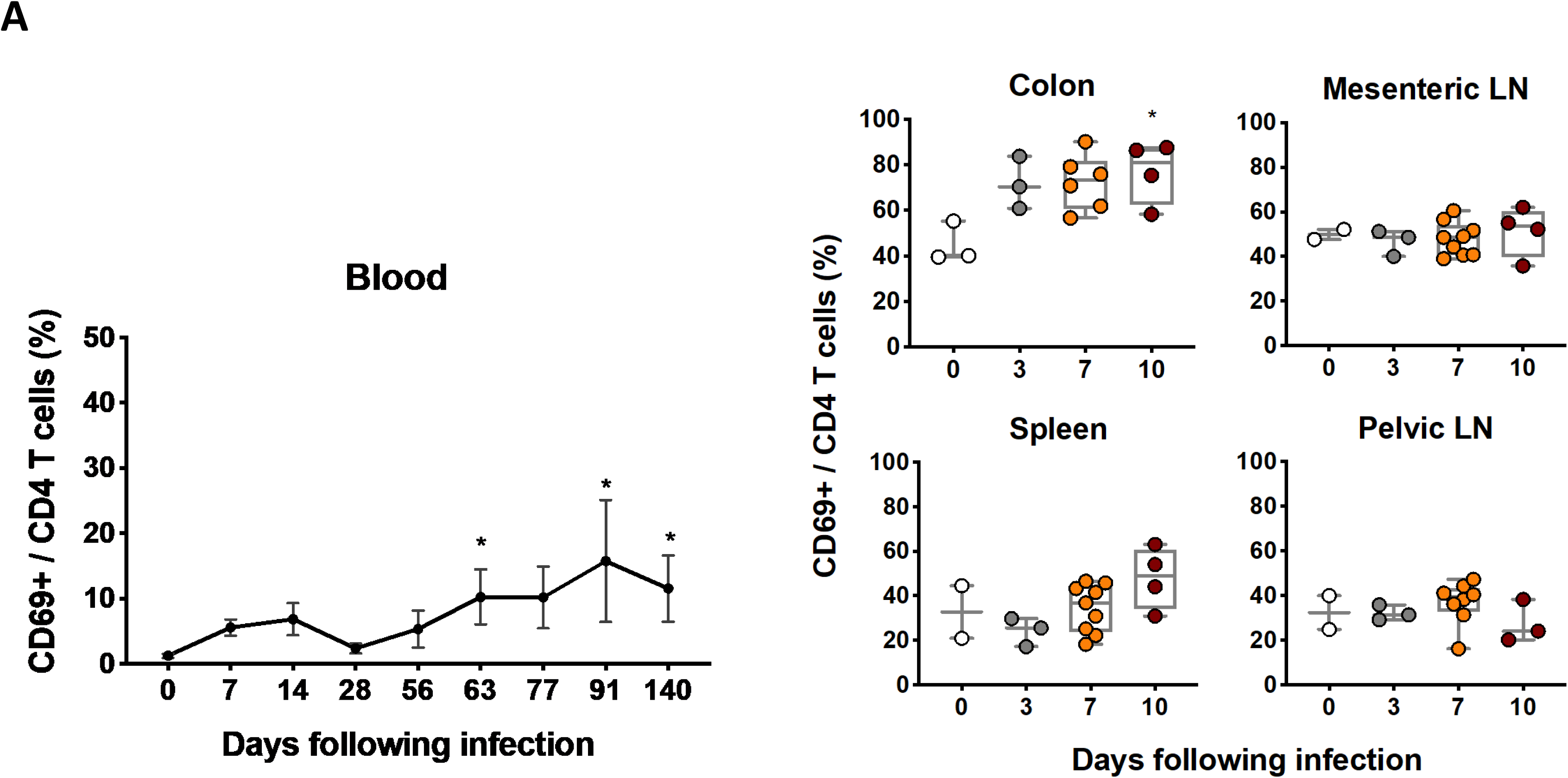
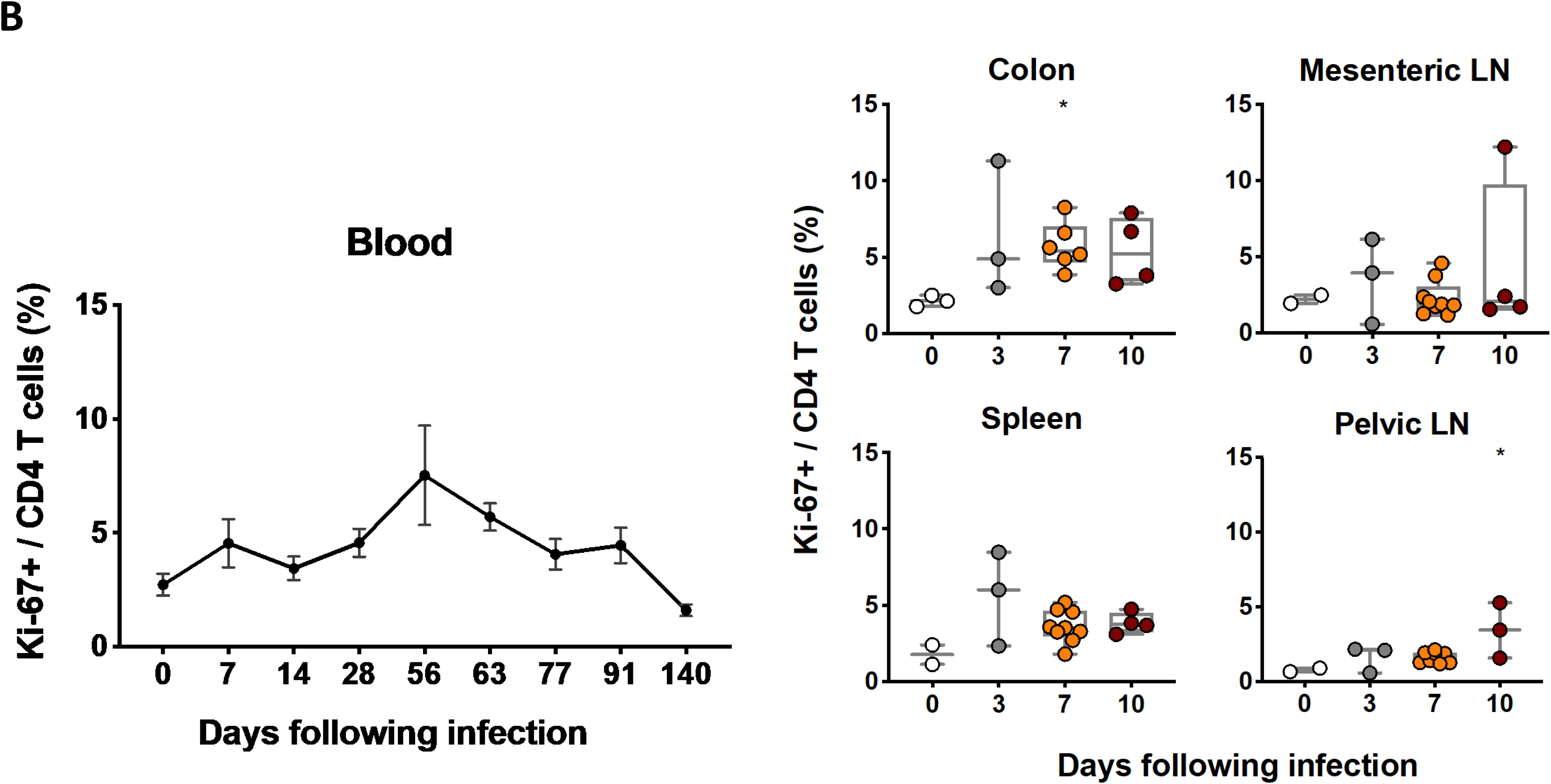
The expression of CD69 and Ki67 in CD4 T cells from blood and tissues. (**A**) CD69 expression of CD4 T cells from blood and multiple tissues at necropsy during the course of SIVmac251 infection ranging from day 0 to day 140 following SIVmac251 infection. (**B**) Ki67 expression of CD4 T cells in blood and multiple tissues at necropsy in monkeys with SIVmac251 infection. Significant *p* values are shown in figures. Experimental variables were analyzed by one-way analysis of variance (ANOVA). *\* p .05*

**Figure S3.**
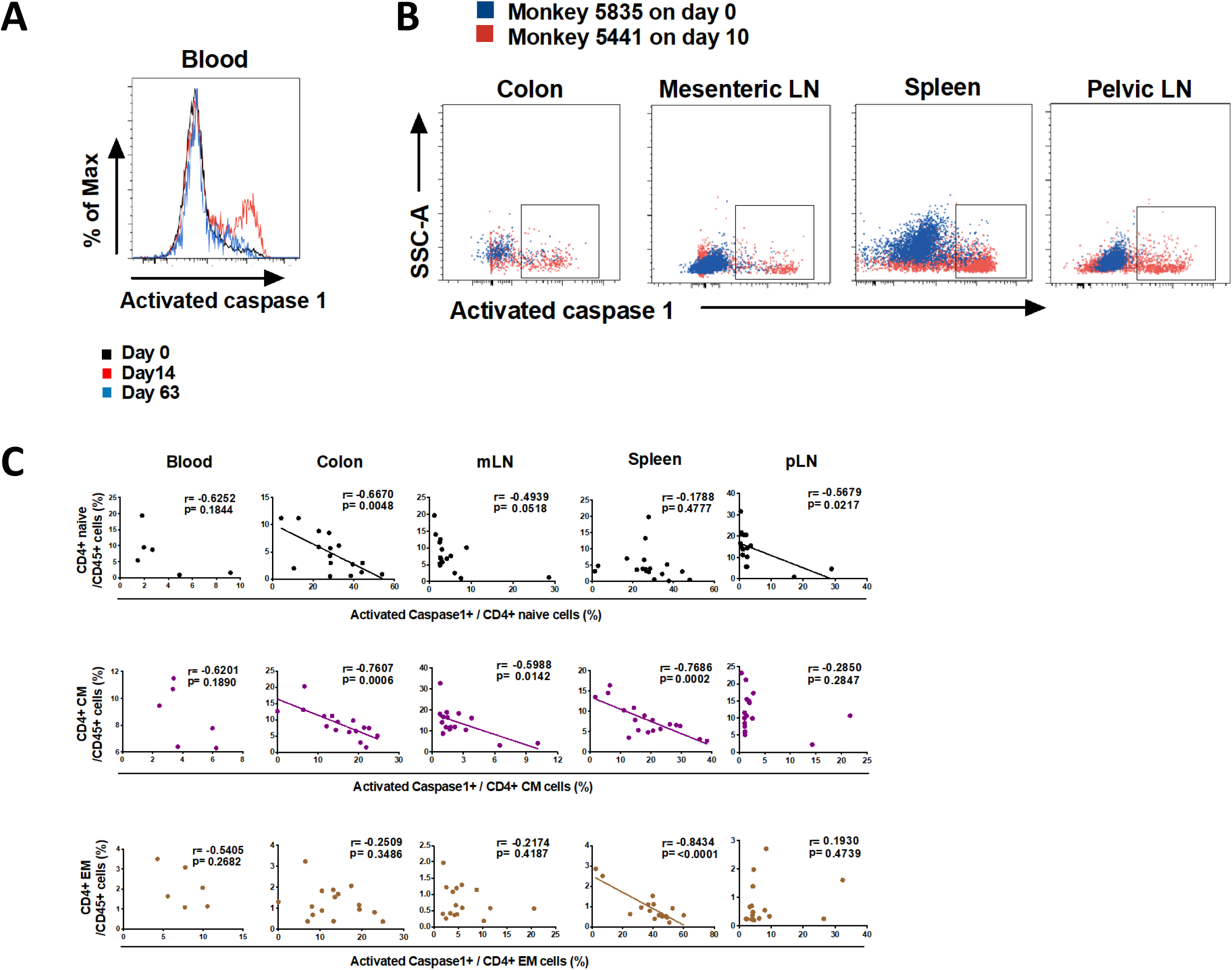
Detection of CD4 T cell pyroptosis during SIV infection. (**A**) Representative flow cytometric histogram indicating pyroptosis of CD4 T cells in blood from one monkey on days 0, 14 and 63. (**B**) Representative flow cytometric plots showing CD4 T cells expressing activated caspase 1 in multiple tissues from two monkeys with or without SIV infection. (**C**) Correlation between the expression of activated caspase 1 in CD4 T cell subsets including naive, central memory (CM), effector memory (EM) subsets and their corresponding level in blood and tissues. Solid lines indicate the correlation with significant p values. spearman rank order correlation coefficients r and corresponding p values are indicated.

**Figure S4.**
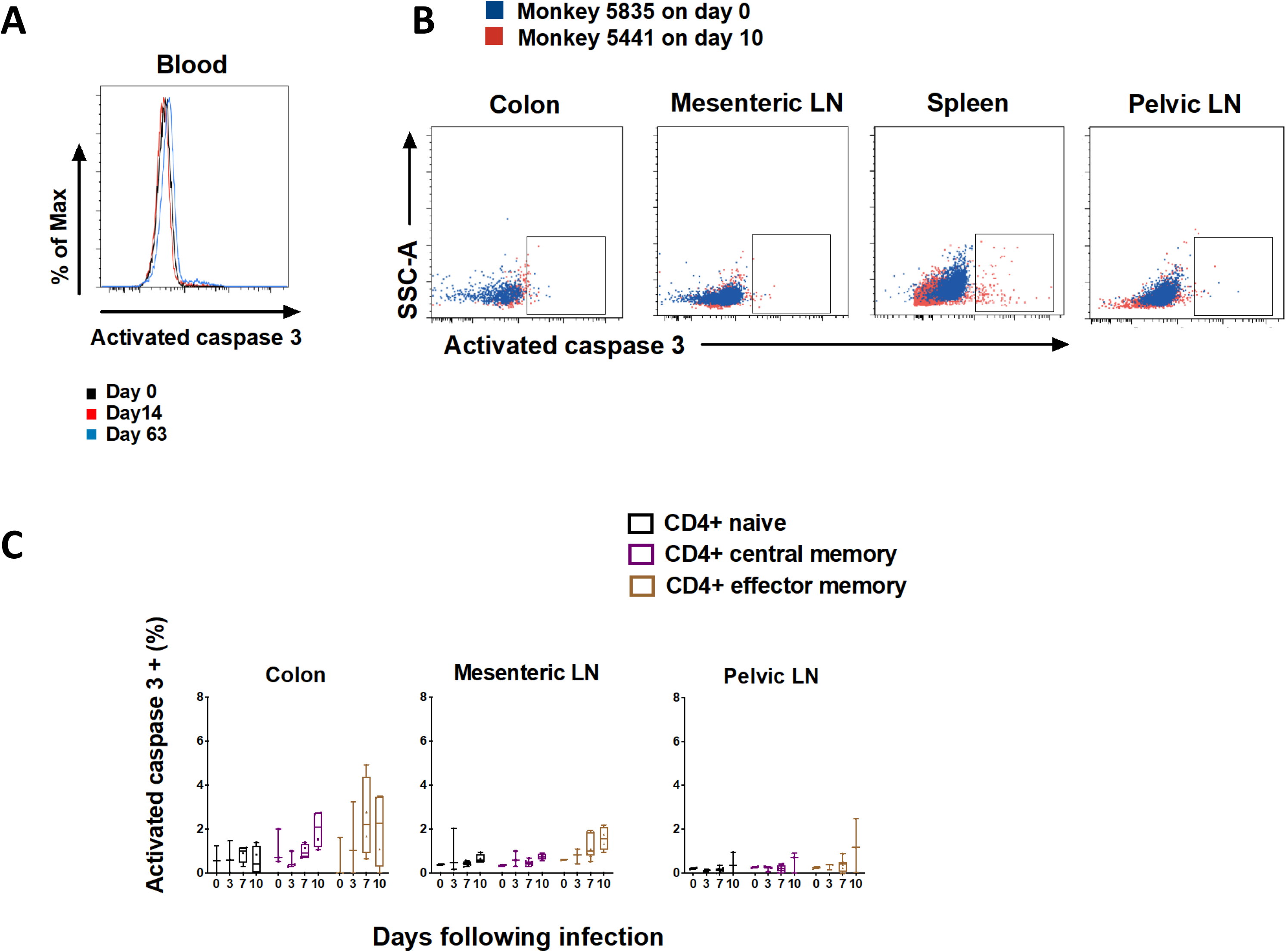
Detection of CD4 T cell apoptosis during SIV infection. (**A**) Representative flow cytometric histogram indicating apoptosis of CD4 T cells in blood from one monkey on days 0, 14 and 63. (**B**) Representative flow cytometric plots showing CD4 T cells expressing activated caspase 3 in multiple tissues from two monkeys with or without SIV infection. (**C**) The apoptosis of CD4 T-cell naïve, central memory and effector memory subsets in colon, mesenteric or pelvic LNs during pathogenic SIVmac251 infection. Black, purple and brown correspond to naive, CM and EM CD4 T-cell subset, respectively.

**Figure S5.**
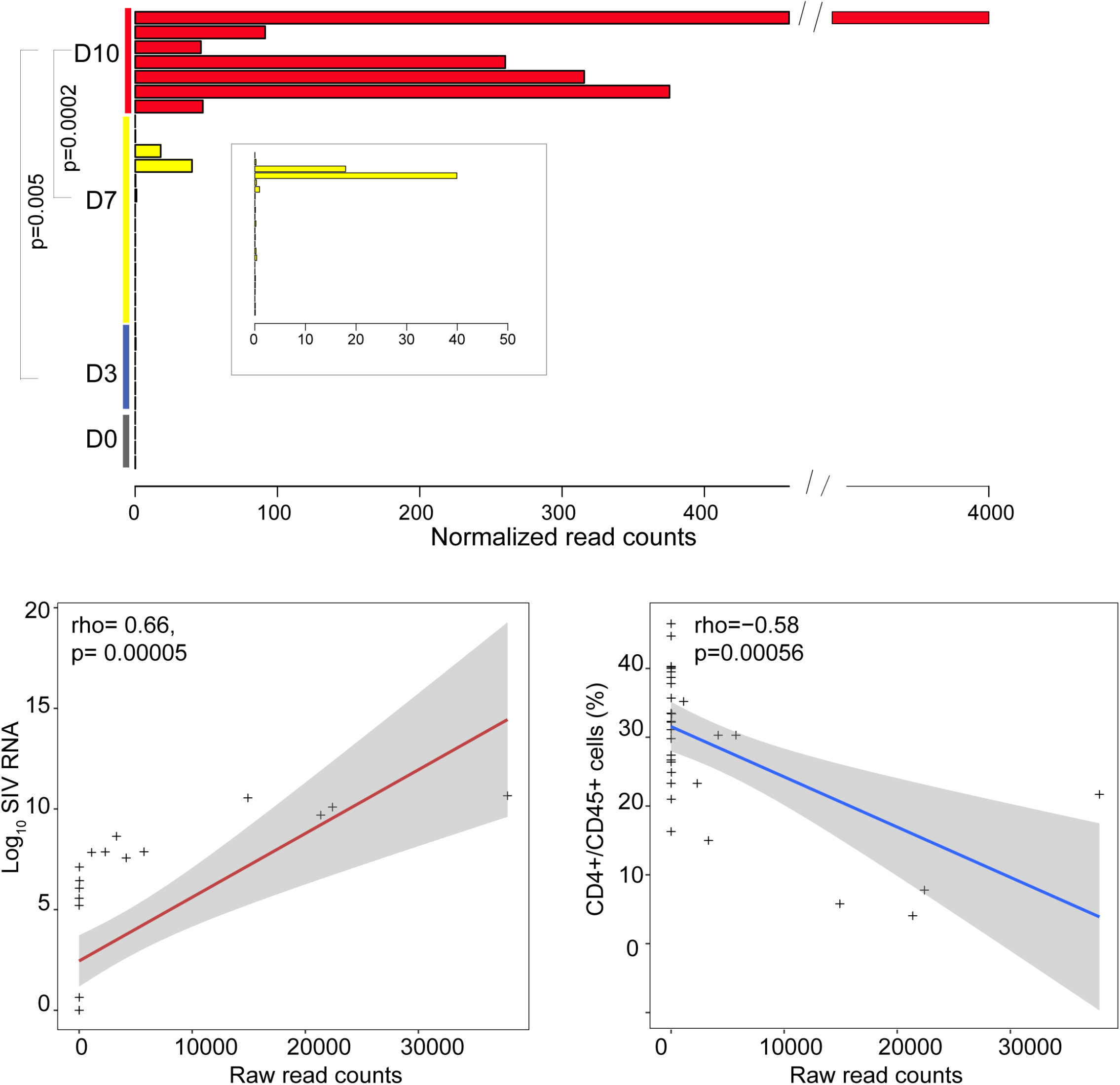
Distribution of RNA-Seq reads that were mapped to the SIV genome at days 0, 3, 7 and 10. At the top, reads that did not map the rhesus macaque genome were mapped to the SIV genome. Counts per millions reads was plotted for each sample at each time point. Enrichment of SIV mapped reads at day 10 compared to day 3 and day 7 was assessed using Wilcox rank test. At the bottom, spearman correlation between the number of reads mapped to the SIV genome for each animal and SIV RNA or the frequency of CD4+ cells. Each dot on the plot represents an animal. Gray are represents the plot interval confidence (IC) of 95%.

**Figure S6.**
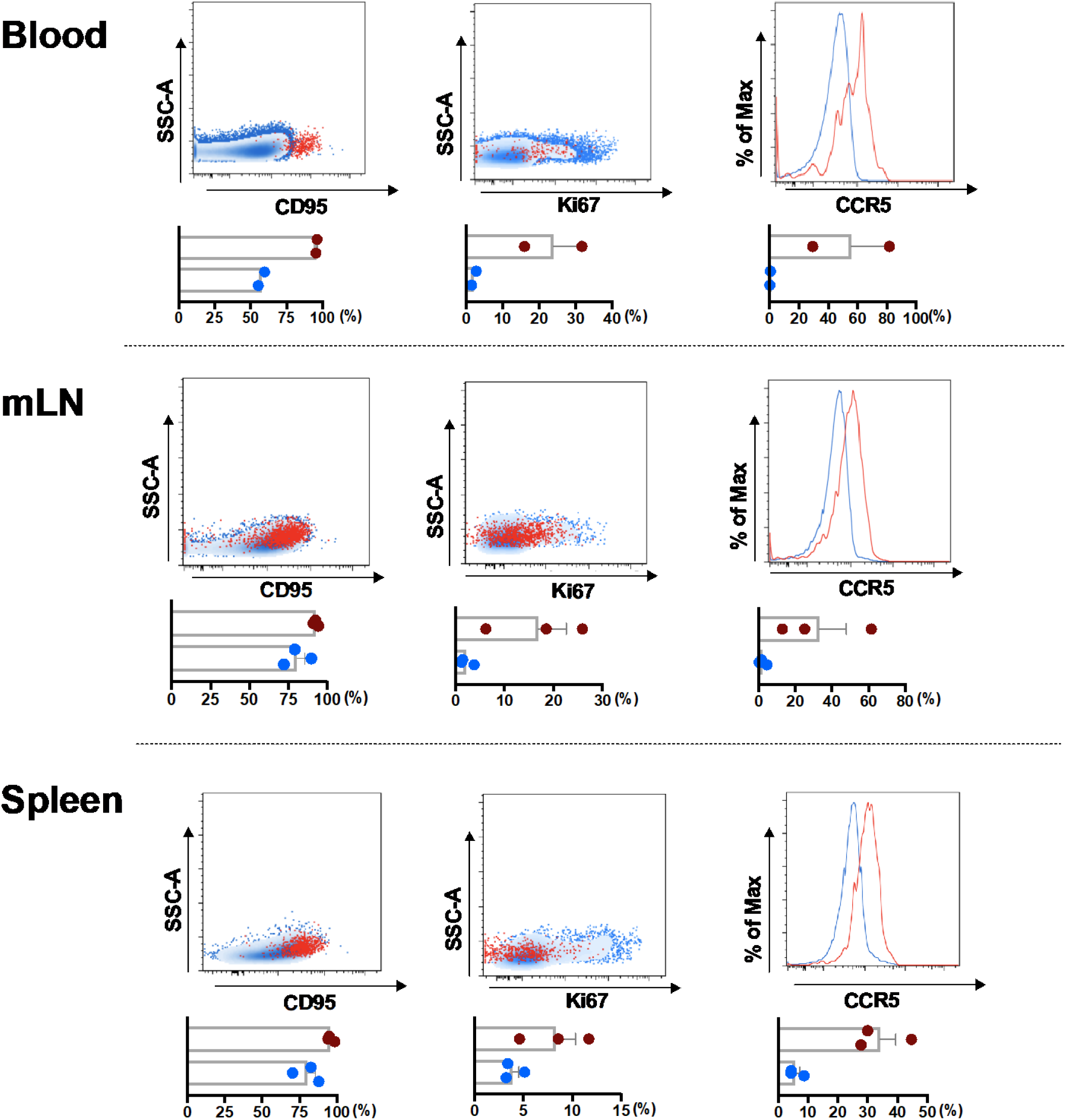
Characterization of CD4 T cells with translation-competent SIV in blood, spleen and mesenteric LNs from SIV-infected monkeys during early SIV infection. From left to right, representative flow cytometric plots or histogram showing CD4 T cells expression CD95, ki67 and CCR5, respectively. Uninfected CD4 T cells are shown in blue, SIV+CD4 T cells are shown in red. The corresponding data are also summarized in column below.

**Figure S7.**
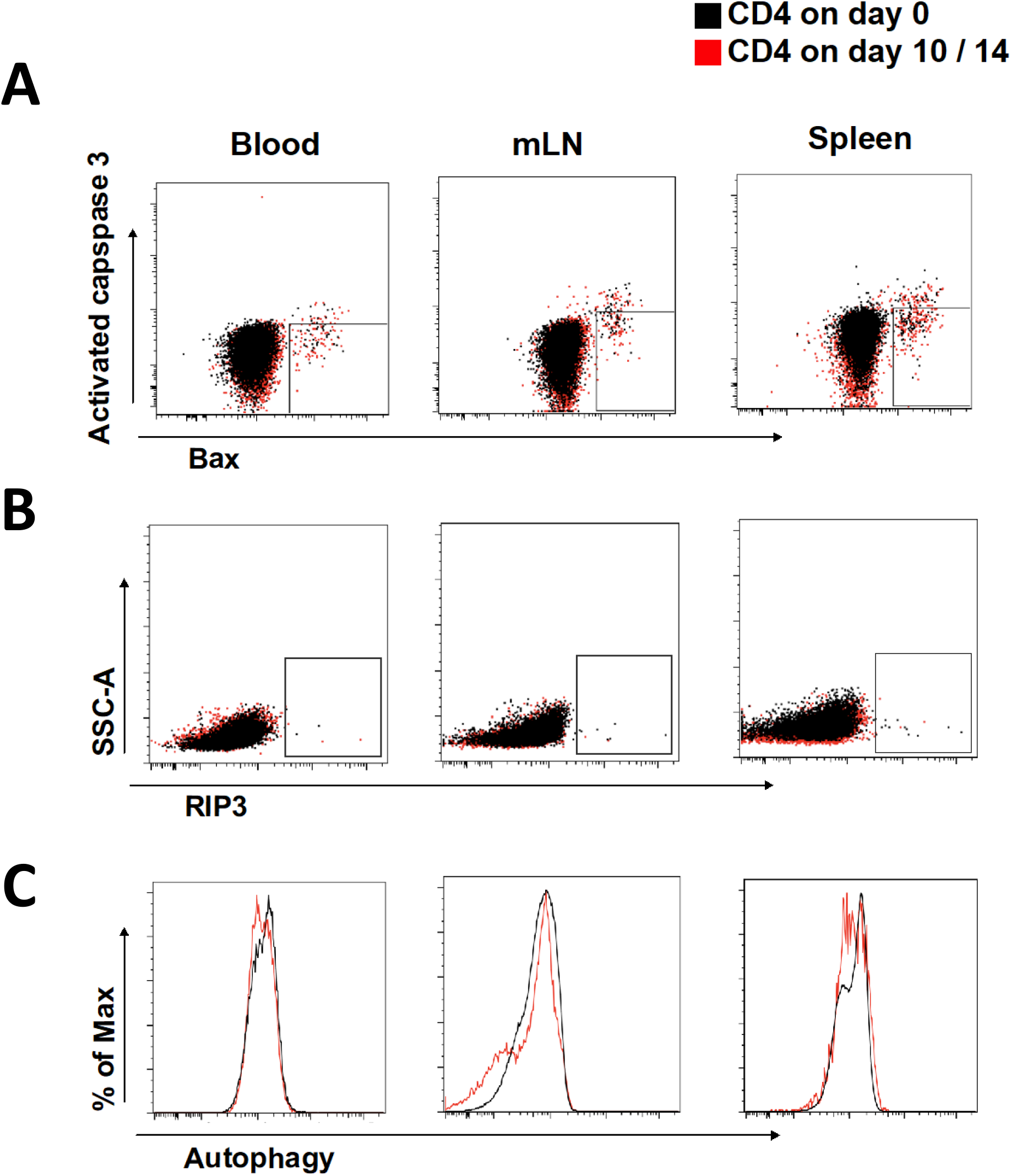
CD4 T cells capable of undergoing mitochondria-induced caspase-independent cell death, necroptosis or autophagy in blood and tissues. (**A**) Representative flow cytometric plots showing CD4 T cells containing Bax but without caspase 3 expression in blood and tissues from naive or infected monkeys during early SIV infection. (**B**) Representative flow cytometric plots showing CD4 T cells expressing RIP3 in blood and tissues from naive or infected monkeys during early SIV infection. (**C**) Representative flow cytometric plots showing CD4 T cells staining with autophagy probes in blood and tissues from naive or infected monkeys during early SIV infection.

**Figure S8.**
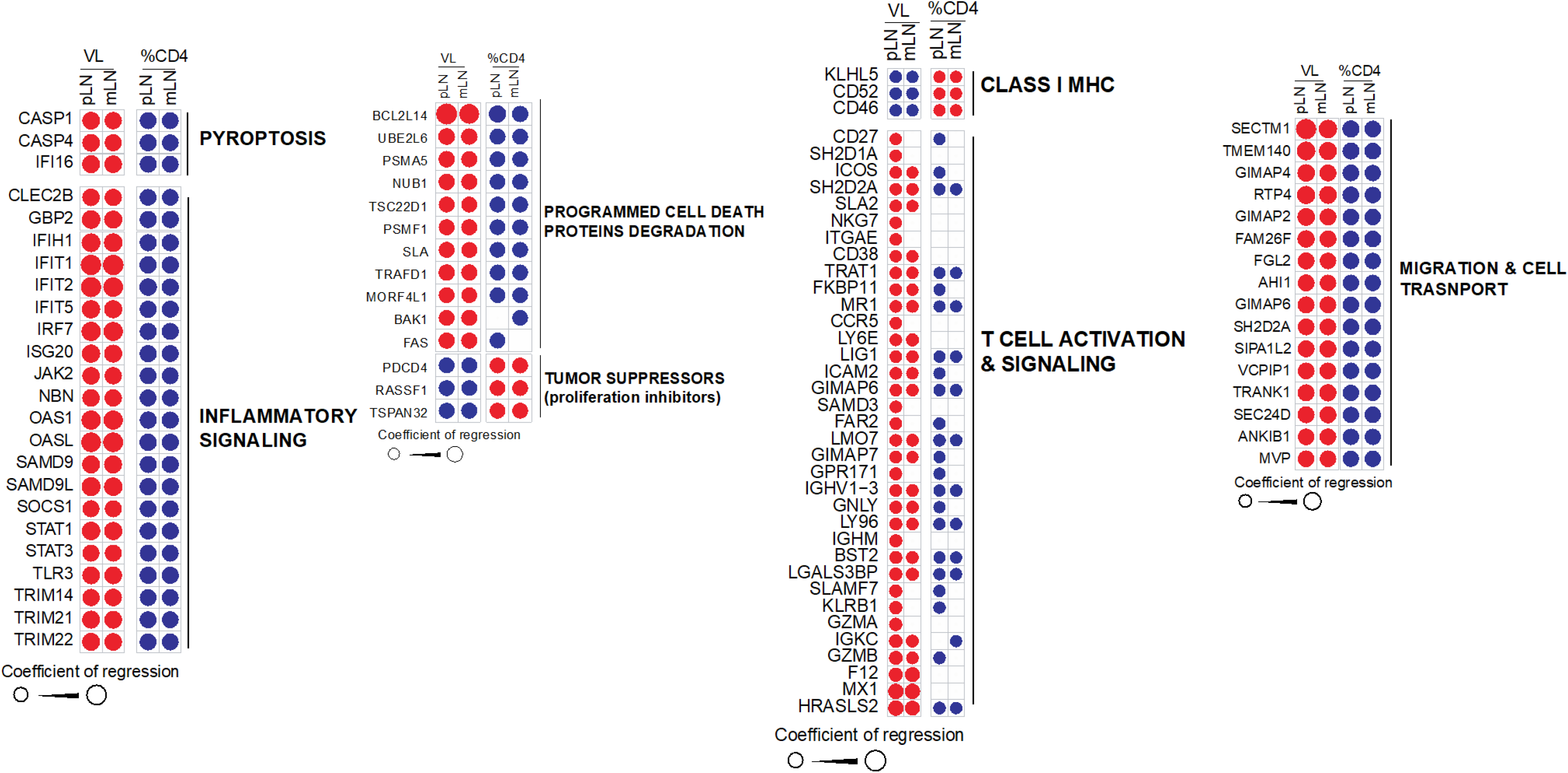
Markers of pyroptosis, inflammatory response, programmed cell death and protein degradation, tumor suppressors, T cell activation and signaling, class MHC I, cell migration and transport that correlated significantly with viral RNA and the frequency of CD4 T cells in pLN and mLN. Red and blue circles show the regression coefficient of the expression of each marker significantly correlated (p < 0.05) with viral RNA or the frequency of CD4 T cells, respectively. Red stands for positive correlation and blue stands for negative correlation. The size of the circle represents the level of the correlation where bigger the size of the circle, higher is the correlation of this gene with viral RNA or frequency of CD4+ cells.

## Methods

### Animals and infections

26 outbred, Indian-origin adult or juvenile female or male rhesus monkeys (Macaca mulatta) were genotyped and selected as negative for the protective MHC class I alleles Mamu-A*01, Mamu-B*08, and Mamu-B*17. TRIM5 polymorphisms were balanced equally among groups, and animals were otherwise randomly allocated. All monkeys were housed at Bioqual, Rockville, MD. 19 animals were infected with 5 × 10^4^ TCID50 of our SIVmac251 challenge stock by the intravaginal route (14), and were necropsied at time points day 0, 3, 7 and 10 following infection. At necropsy, multiple tissues including colon, spleen, mesenteric lymph nodes and pelvic lymph nodes were obtained and processed shortly after collection. 7 monkeys were infected with SIVmac251 challenge stock by intrarectal route, and blood were collected at multiple time points from day 0 to day 140. All animal studies were approved by the appropriate Institutional Animal Care and Use Committee (IACUC).

## METHOD DETAILS

### Viral RNA and DNA

Plasma SIV RNA levels were detected using a *gag*-targeted quantitative real-time RT-PCR assay, and levels of SIV RNA or DNA in tissues were measured using *gag* targeted, nested quantitative hybrid real-time/digital RT-PCR and PCR assays as previously described (14).

### Whole-blood lymphocyte count assay

100 ul whole blood were stained with monoclonal antibodies CD3 (V450 conjugate), CD4 (PE conjugate) and CD8 (FITC conjugate), then lysed in TQ-Prep and fixed. The acquisitions were performed on LSRFortessa and flow cytometric analysis was performed by FlowJo software to generate the percentage of CD4 T cells within lymphocytes. 800 ul whole blood was obtained from animals and Compete Blood Count (CBC) performed on Siemens Advia 120. CD4 T-cell count was determined by combining cell counts of lymphocytes with the frequency results generated by flow cytometry.

### Flow cytometric analysis

The LIVE/DEAD™ Fixable Aqua Dead Cell Stain Kit was applied to excluding the dead cells. Cell surface staining was conducted using the following monoclonal antibodies including CD45 (BV605 conjugate, clone D058-1283, BD Pharmingen), CD3 (Alexa 700 conjugate, clone SP34.2, BD Pharmingen), CD4 (BB700 conjugate, clone L200, BD Pharmingen), CD8 (Pacific Blue conjugate, clone RPA-T8, BD Pharmingen), CD95 (BV711 conjugate, clone DX2, BD Pharmingen), CD28 (PE-Cy7 conjugate, clone 28.2, ThermoFisher), CD69 (ECD conjugate, clone TP1.55.3, Beckman Coulter). For intracellular staining, cells were permeabilized using Caltag Fix & Perm (Invitrogen), then stained with monoclonal antibodies including Ki67 (PE conjugate, clone B56, BD Pharmingen) and caspase 3 (BUV395 conjugate, clone C92-605, BD Pharmingen).

Fluorescent labelled inhibitor probe FAM-YVAD-FMK (ImmunoChemistry Technologies) which could covalently bind to the activated caspase 1, was applied to detecting the cells containing activate caspase 1. Autophagy probe (ImmunoChemistry Technologies) which fluoresces red when inserting into the lipid membrane of autophagosomes and autolysosomes was applied in the study. After staining, cells were washed and fixed by 2% paraformaldehyde. All data were acquired on BD LSRII flow cytometer and analyzed by FlowJo software (version 9.9.5).

### Transcriptomic Analysis

CD4 T cells from mesenteric LNs and pelvic LNs in monkeys necropsied on day 0, 3, 7 and 10 following infection were enriched by negative selection through EasySep™ Non-Human Primate Custom Enrichment Kit (STEMCELL Technologies). The enriched cells were further sorted by BD FACSAria flow cytometer. RNA extraction was carried out using RNeasy Mini Kit (QIAGEN) according to manufacturer’s instructions. A Low-Input mRNA library (Clontech SMARTer) v4 was applied and samples were sequenced on Illumina NS500 Paired-End 75 bp (PE75) at the Molecular Biology Core Facility at Dana-Farber Cancer Institute. All samples were processed using an RNA-Seq pipeline implemented in the bcbio-nextgen project (https://bcbio-nextgen.readthedocs.org/en/latest/). Raw reads were examined for quality issues using FastQC (http://www.bioinformatics.babraham.ac.uk/projects/fastqc/) to ensure library generation and sequencing are suitable for further analysis. Adapter sequences, other contaminant sequences such as polyA tails and low-quality sequences with PHRED quality scores less than five were trimmed from reads using atropos (https://github.com/jdidion/atropos). Trimmed reads were aligned to the Macaca mulatta genome (https://www.unmc.edu/rhesusgenechip/index.htm) using STAR v2.7.3. Alignments will be checked for evenness of coverage, rRNA content, genomic context of alignments (for example, alignments in known transcripts and introns), complexity and other quality checks using a combination of FastQC, Qualimap and other custom tools. Counts of reads aligning to known genes are generated by featureCounts. We used DESeq2 (Moderated estimation of fold change and dispersion for RNA-Seq data with DESeq2) internal filtering algorithm called “independentFiltering” that sets a threshold on the mean of normalized counts of all samples (baseMean > 10) in order to maximize the number of tests that pass multiple test correction. A corrected P value cut-off of 0.05 was used to assess significant genes that were up-regulated or down-regulated on days 3, 7, and 10 using an adjusted P of<0.05 and the Benjamini-Hochberg (BH) correction method. Gene set enrichment analysis (30, 31) and database of biological and immunological modules (https://www.gsea-msigdb.org/gsea/index.jsp) were used to explore enriched pathways at days 3, 7 and 10 after infection. An FDR q value cut-off of 0.05 was used to selected increased or decreased pathways at days 3, 7 and 10 compared to day 0. Raw fastq files were uploaded to NCBI Gene Expression Omnibus (GEO) database under identifier GSE165519.

### Mapping of SIV RNA

In order to test for the presence of SIV RNA at days 3, 7 and 10 following SIVmac251 infection, unmapped reads, that did not align to the rhesus genome by STAR aligner were extracted and mapped to the SIVmac251 genome using Burrows-Wheeler Aligner bwa mem (https://github.com/lh3/bwa).

### SIV RNA-Flow FISH

Prior to SIV mRNA detection, cells were stained with viability dye, then surface staining were performed to identify phenotype of cell populations using following monoclonal antibodies including CD45 (BV786 conjugate), CD4 (BB700 conjugate,), CD3 (BV421 conjugate), CD195 (BV650 conjugate), CD8 (APC-Cy7 conjugate), CD95 (BUV737 conjugate), followed by intracellular staining with SIV Gag protein p27 (FITC conjugate, clone 55-2F12, conjugated in house) and Ki67 (PE-Cy7 conjugate). PrimeFlow RNA assay was carried out according to manufacturer’s instructions (Thermo Fisher Scientific). Generally, The SIV specific oligonucleotide target probe set that contains 22∼50 pairs could bind to SIV *env, gag* and *pol* mRNA, then signal amplification is achieved through sequential hybridization with multiple amplifier molecules. All data were acquired on BD LSRII flow cytometer and analyzed by FlowJo software (version 9.9.5).

### Statistical analysis

All the statistical and graphic analyses were done using GraphPad Prism software (version 7.0). For comparisons between groups, Kruskal–Wallis one-way analysis of variance was applied in the study. Spearman’s rank correlation was used to analyze the correlation between variables. P values less than 0.05 were considered to be significant in the study.

